# *Shank3* mutations impair electrical synapse scaffolding and transmission in mouse brain

**DOI:** 10.1101/2021.03.25.437056

**Authors:** Jonathan Lautz, Zhiyi Zhu, Haley E. Speed, Stephen E. P. Smith, John P. Welsh

## Abstract

*Shank3* mutations contribute to intellectual disability. Because SHANK3 is a protein scaffold that helps organize the multiprotein network of the glutamatergic postsynaptic density (PSD), alterations in chemical synaptic transmission are implicated. Electrical synaptic transmission is a second form of synaptic transmission, enabled by intercellular channels comprised of connexin36 that support direct electrical communication among neurons, electrical brain rhythms, and neurocognitive states. Using multiplex proteomics, we report that two autism-related mutations of mouse *Shank3* disrupt the glutamatergic PSD differently, but have in common the disruption of an association between NMDA-type glutamate-receptors (NMDARs) and connexin36. Mutation of *Shank3* exons 13-16 most robustly dissociated connexin36 from NMDARs while impairing electrical synaptic transmission and the synchrony of an electrical rhythm in mouse inferior olive. We suggest that electrical synapses are a component of an “extended PSD” sensitive to *Shank3* mutations that produce intellectual disability, at least in part, by impairing electrical synaptic transmission.

## INTRODUCTION

Mutations of *Shank3* produce intellectual disability (ID; Wilson et al., 2003) that can be comorbid with schizophrenia (Gauthier et al., 2010) and autism spectrum disorder (ASD, Betancur and Buxbaum, 2013; Grabrucker, 2014; Sala et al., 2015; Uchino and Waga, 2013). SHANK3 is a scaffold protein that helps organize the multiprotein network comprising the postsynaptic density (PSD) of glutamatergic synapses due to its five binding domains (Naisbitt et al., 1999). By its PSD-drosophila disc large tumor suppressor-zonula occludens 1 (PDZ) domain, SHANK3 binds AMPA- and NMDA-type glutamate receptor subunits either directly or *via* other PDZ-bound scaffold proteins such as guanylate-kinase-associated protein (GKAP), PSD protein-95 (PSD-95), and SAP90/PSD-95 associated protein (SAPAP) (Boeckers et al., 1999; Monteiro and Feng, 2017; Ponna et al., 2018; Uchino et al., 2006). The SHANK3 PDZ domain also binds the signaling effectors Ca^2+^/calmodulin-dependent protein kinase II (CaMKII) and synaptic Ras-GTPase-activating protein (SynGAP; Tao-Cheng et al., 2014; Walkup et al., 2016). By its C-terminus proline-rich region and sterile α motif domain, SHANK3 binds the scaffold protein Homer, forming a mesh-like matrix (Tu et al., 1999; Hayashi et al., 2009; Kursula, 2019). Homer, in turn binds to group 1 metabotropic glutamate receptors (mGluRs; Mao et al., 2005), transient receptor potential canonical family channels (Smani et al., 2014), and cytoskeletal elements such as dynamins (Gray et al., 2003).

There is general agreement that ID with *Shank3* mutations results from impairments in glutamatergic synaptic transmission due to the improper integration of signaling and receptor proteins within the PSD. *Shank3* transcripts are expressed ubiquitously and are enriched in layer II/III neocortex, hippocampus, striatum, thalamus, cerebellum, and the inferior olive (IO) (Lein et al., 2007) where plasticity of glutamatergic synapses encodes experience. However, the contribution of any impairment in neuronal communication other than by chemical synaptic transmission has not been addressed.

Electrical synaptic transmission is a second form of synaptic transmission whose relation to ID has not been examined. Electrical synapses between brain neurons are established by plaques of intercellular channels (gap junctions) constructed of the protein connexin36 (Cx36) (Condorelli et al., 1998; Faber and Pereda, 2018). Electrical synapses allow current to flow between neurons and have an advantage over chemical synapses in the speed and continuity of information transfer, because direct electrical communication is not time-limited by axonal conduction and neurotransmitter diffusion. Electrical synaptic transmission is important because it can synchronize spike output among neuronal ensembles to drive coherent brain activity in distant postsynaptic targets, such as occurs in the flow of activity from the thalamus to the neocortex (Coulon and Landisman, 2017) and from the IO to the cerebellum (Llinás et al., 1974; Llinás and Sasaki, 1989; Welsh et al., 1995; Welsh and Llinás, 1997; Welsh and Turecek, 2017). Like SHANK3, electrical synapses are expressed throughout the brain.

We combined multiplex proteomics and measurements of electrical coupling to test whether SHANK3 integrates Cx36 into the PSD multiprotein network to influence the synchrony of a brain rhythm. The studies were informed by studies of the IO demonstrating the proximity of gap junctions to the glutamatergic PSD (Hoge et al., 2011; Turecek et al., 2014) where electrical synaptic transmission is highly prevalent (Devor and Yarom, 2002), making the IO an ideal structure in which to study the contribution of electrical synapses to the intrinsic oscillatory properties of the brain (Llinás, 2014). The studies were further motivated by the identification of a PDZ binding domain on the C-terminus of Cx36, a candidate for SHANK3 binding (Li et al., 2004). We examined two mouse lines with *Shank3* mutations: one heterozygous for lacking *Shank3* exon 21 (Shank3C mutation) which deletes the C-terminus Homer binding sites (*Shank3^+/ΔC^* or HetC mice; Kouser et al., 2013) and a second lacking *Shank3* exons 13-16 (Shank3B mutation) encoding the PDZ binding domain (*Shank3^B/B^* or HomB mice; *Shank3^+/B^* or HetB mice; Peça et al., 2011). Mutations in both regions of *Shank3* have been implicated in human ID (Li et al., 2018; Zhu et al., 2018). Mice heterozygous for the Shank3C mutation (Kloth et al., 2015) and mice heterozygous or homozygous for the Shank3B mutation (Jaramillo et al., 2017; Peça et al., 2011) exhibited behavioral phenotypes consistent with ASD and obsessive compulsive disorder.

We report a structural association of Cx36 to the NMDAR within the glutamatergic PSD of IO neurons that is disrupted by two *Shank3* mutations, most significantly by deleting the SHANK3 PDZ binding domain which impairs electrical synaptic transmission, neural network synchrony, and the strengthening of both by NMDAR activation. The work emphasizes that the glutamatergic PSD and neuronal electrical synapses are integrated in the mammalian brain and that impairments in electrical synaptic transmission are likely to contribute to syndromes due to *Shank3* mutation.

## RESULTS

### Cx36 is enriched in the glutamatergic PSD

We first tested whether Cx36 is enriched in the PSD subcellular fraction of brainstem homogenates containing the IO. A previous study demonstrated enrichment of Cx36 in the PSD subcellular fraction of whole rat brain homogenate (Lynn et al., 2001). PSD fractions comparing the relative amounts of Cx36 and NR1 subunit of the NMDAR (NMDAR1) were analyzed by Western immunoblot (Figure 1A). PSD-95 and NMDAR1 were enriched in PSD fractions, as expected (Figure 1B,C). Cx36 was enriched in PSD fractions also, indicating a possible structural basis for the association of neuronal gap junctions to the glutamatergic PSD previously shown by electron microscopy (Hoge et al., 2011). HetC mice showed significant reduction of NMDAR1 in the PSD fraction relative to WT (73 ± 21% of WT; *p* = 0.03), indicating reduced membrane insertion of the NMDAR due to SHANK3 C-terminus deletion (Figure 1D). This effect was not observed in HomB mice (103 ± 5% of WT; *p* = 0.6; Figure 1E), indicating that deleting the SHANK3 PDZ binding domain did not affect PSD expression of NMDAR1.

**Figure 1.**
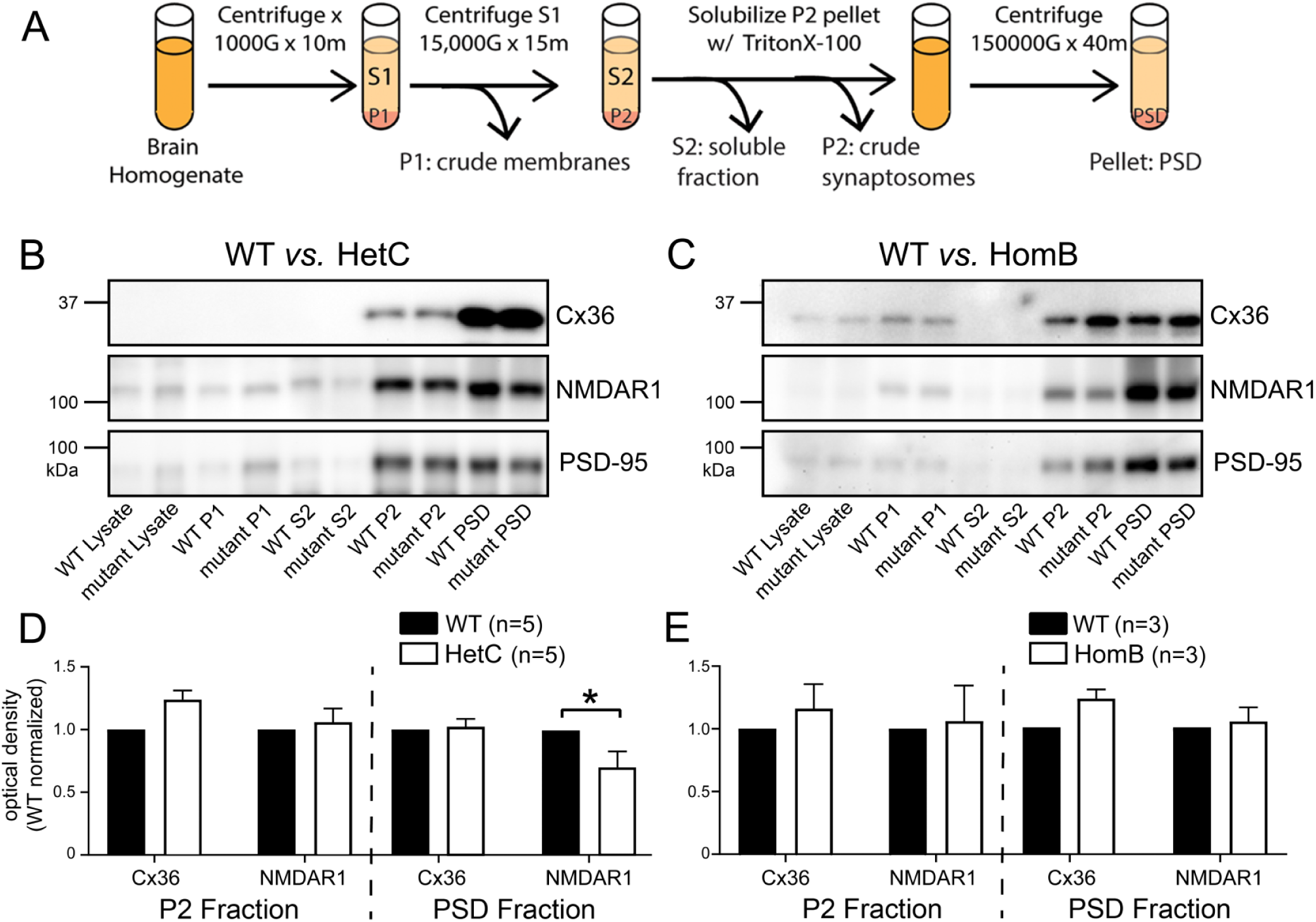
Cx36 is enriched in the glutamatergic PSD. (A) Protocol for PSD preparation. (B,C) Western blots showing relative levels of Cx36, NMDAR1, and PSD-95 in subcellular fractions of IO tissue from HetC, HomB, and WT mice. Lanes: lysate, tissue homogenate; P1, nuclear pellet; S2, cytosolic fraction; P2, synaptosomal pellet; PSD, PSD pellet. Blots were probed for the protein indicated in each set of lanes. (D,E) Mean WT-normalized blot density for the P2 and PSD fractions from HetC and HomB mice (* *p* < 0.05, paired t-test). *N* = 16 mice.

### SHANK3 PDZ binding associates Cx36 with NMDAR1

We next tested by co-immunoprecipitation (co-IP) whether a structural association of Cx36 to NMDAR1 contributed to the enrichment of Cx36 within the PSD. Cx36 and NMDAR1 were immunoprecipitated from IO dissections of WT, HetC, and HomB mice and the relative amounts of co-IP’d Cx36 and NMDAR1 were analyzed by Western immunoblot (Figure 2). Cx36 and NMDAR1 showed robust associations in WT mice when either Cx36 or NMDAR1 was the precipitant (Figure 2A,B, WT). This was the first direct indication that chemical and electrical synaptic transmissions in the mammalian brain are structurally related. HetC mice did not differ from WT in the co-IP of Cx36 or NMDAR1 (5 ± 11% change, *p* = 0.5) (Figure 2A,C). In contrast, HomB mice exhibited a significant 31 ± 11% decrease in the Cx36-NMDAR1 association (*p* = 0.02) when Cx36 was immunoprecipitated, providing evidence that the two were associated in a multiprotein complex that required the SHANK3 PDZ binding domain (Figure 2B,D). Of note, HomB mice exhibited significant reduction in the co-IP of NMDAR1, indicating either reduced NMDAR1 co-association within the PSD or reduced overall expression.

**Figure 2.**
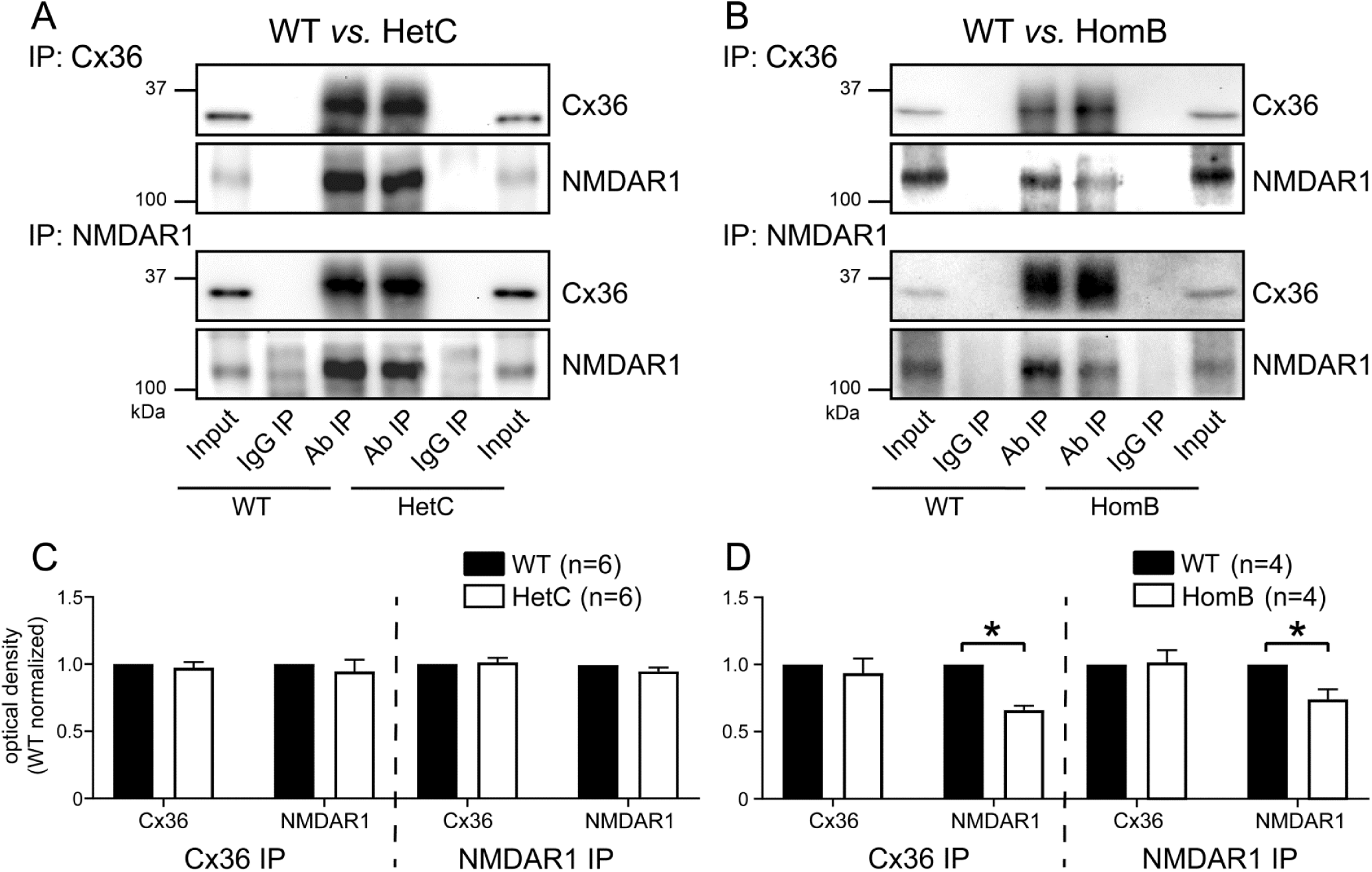
SHANK3 PDZ binding associates Cx36 with NMDAR1. (A,B) Co-IP Western blots showing relative amounts of Cx36 and NMDAR1 in IO tissue from HetC and HomB mice compared to WT. Cx36 or NMDAR1 was IP’d and the blots probed for the protein indicated in each set of lanes. Lanes: Input, tissue lysate without IP; IgG IP, lysate immunoprecipitated with non-specific IgG; Ab IP, lysate immunoprecipitated with IP antibodies. (C,D) Mean optical density normalized to WT showing reduced Cx36-NMDAR1 association in HomB mice when Cx36 is the precipitant (* *p* < 0.05, t-test). *N* = 20 mice.

### SHANK3 scaffolds Cx36 into the synaptic multiprotein network

We examined how two *Shank3* mutations affect the relationship of Cx36 to the synaptic multiprotein network using quantitative multiplex co-IP (QMI; Figure 3A). QMI quantifies proteins in shared complexes detected via exposed surface epitopes (PiSCES) using panels of IP and probe antibodies conjugated to Luminex beads and phycoerythrin (PE), respectively, and measured on a modified flow cytometer (Smith et al., 2016, Lautz et al., 2018, 2019). We used QMI to examine changes in 418 PiSCES using a panel of validated antibodies (Lautz et al., 2018) recognizing Cx36 and 20 PSD proteins (6 receptor proteins, 6 signaling proteins, 8 structural proteins; Supplemental Information).

**Figure 3.**
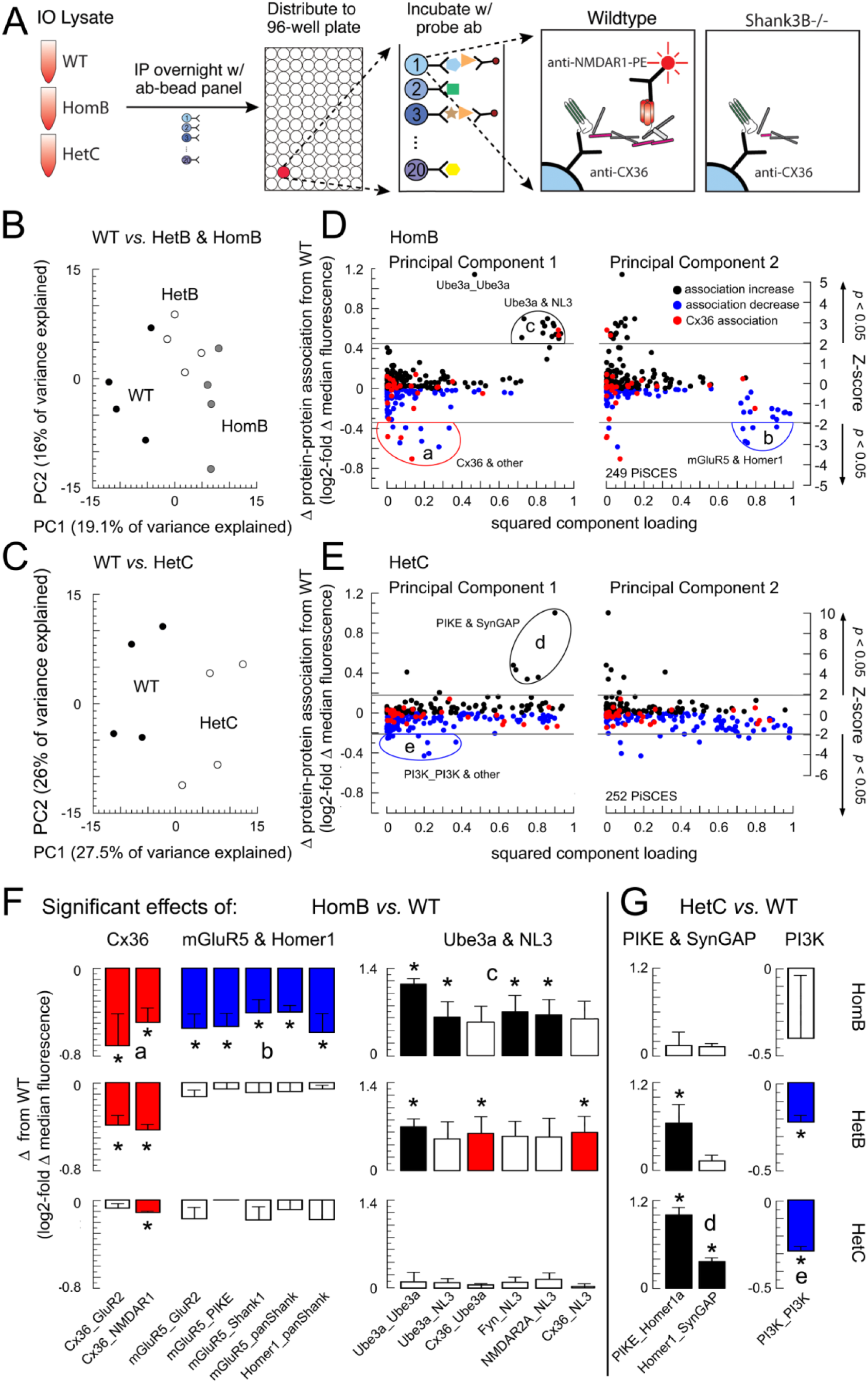
Synaptic protein associations disturbed by *Shank3* mutations include Cx36. (A) QMI methods. IO lysates from WT and *Shank3* mutant mice were incubated with a panel of antibody-coupled latex microspheres, which were distributed into wells of a 96-well plate and incubated with 21 PE-coupled probe antibodies (1 antibody per well) to detect 400+ binary PiSCES. After PE detection using a refrigerated flow cytometer, mutant mice were compared to matched WT controls using customized statistical programs to detect significantly changed PiSCES. (B,C) Separation of Shank3B and Shank3C mutant mice from WT by PCA. Each circle is one mouse. (D,E) The change in magnitude for every measured PiSCES plotted against the squared component loading for the first two principal components. Each dot shows a single PiSCES averaged over 4 mice. Ellipses enclose PiSCES significantly changed by *Shank3* mutation as assessed by Z-score calculated from the mean change and standard deviation of all PiSCES. Horizontal lines show two-tailed, *p* < 0.05 cutoffs. (F,G) PiSCES within elliptical regions in D and E that differed from zero by one-sample t-test (asterisks) for HomB (F) and HetC (G) mice. Bars show mean ± SEM from 4 pairs of mice. * *p* < 0.05. Letters indicate enclosed regions in D and E. Red, PiSCES involving Cx36 with significant change in HomB mice; Blue, PiSCES showing significant decreases in association; Black, PiSCES showing significant increases in association. Changes to PiSCES that are significantly changed in either HomB or HetC mice are demonstrated across genotypes.

#### Principal Component Analysis (PCA)

PCA tested whether *Shank3* genotypes could be distinguished by the change in PiSCES within the IO. In separate PCA analyses, the first two principal components robustly separated HomB, HetB, and WT mice (Figure 3B) and HetC from WT mice (Figure 3C). Follow-up analyses identified subsets of PiSCES in which changes by *Shank3* mutation were responsible for genotype separation.

##### Shank3B

Statistically significant changes in 11 PiSCES due to the Shank3B mutation centered around 4 proteins: 1) Cx36, 2) metabotropic glutamate receptor subunit subtype 5 (mGluR5), 3) ubiquitin protein ligase E3A (Ube3a), and 4) neuroligin 3 (NL3). It was both notable and unexpected that *Shank3* mutations changed multiple PiSCES involving Cx36, implying that SHANK3 links Cx36 to the network of postsynaptic proteins. In HomB mice, the strongest dissociations occurred in PiSCES containing Cx36 and NMDAR1 (expressed as Cx36_NMDAR1, 0.5 ± 0.1 log2-fold, 30% decrease) and Cx36_GluR2 (AMPA-type glutamate receptor subunit 2; 0.7 ± 0.3 log2-fold, 40% decrease) (Figure 3D, region a; Figure 3E, red). Both Cx36 dissociations were observed also in HetB mice but at lower magnitude (0.4 ± 0.1 log2-fold decrease; Fig. 3F, red). All dissociations involving mGluR5 were specific to HomB mice and included mGluR5_GluR2, mGluR5_PIKE (phosphoinositide-3 kinase enhancer), and mGluR5_SHANK (Figure 3D region b; Figure 3F, blue). Significant increases in Ube3a_Ube3a and Ube3a_NL3 (neuroligin-3) also were prominent in HomB mice, as were significant increases in NL3_Fyn (proto-oncogene tyrosine-protein kinase Fyn) and NL3_NMDAR2A (NMDA receptor subunit 2A) (Figure 3D, region c; Figure 3F, black). HomB mice also demonstrated significant dissociation (0.6 ± 0.2 log2-fold decrease; Figure 3F, blue) of Homer1_SHANK (Figure 3D, region b), confirming partial disassembly of the PSD core.

##### Shank3C

PCA separation of HetC and WT mice primarily was due to increases in Homer1a_ PIKE, Homer1a_Homer1, and Homer1a_SynGap (synaptic Ras GTPase-activating protein) (Figure 3E, region d; Figure 3G, black). HetC mice also showed significant dissociations of PI3K_PI3K (phosphoinositide-3 kinase) (Figure 3E, region e; Figure 3G, blue) and a small but statistically significant decrease in Cx36_NMDAR1 (0.1 ± 0.01 log2-fold; 7% decrease; Figure 3F, red). Of note, the only change common to Shank3C and Shank3B mice was the decrease in Cx36_NMDAR1.

#### Adaptive Non-Parametric Analysis with Weighted Cutoff (ANC) and Correlation Network Analysis (CNA)

ANC *∩* CNA analyses (Smith et al., 2016) affirmed that the dissociations of Cx36 from NMDAR1 and GluR2 were among the most prominent consequences of Shank3B mutation (Figure 4A-C). ANC *∩* CNA identified 7 significantly reduced PiSCES in HomB mice: Cx36_NMDAR1, Cx36_GluR2, mGluR5_NMDAR1, mGluR5_PIKE, mGluR5_SHANK, NMDAR2A_Ube3a, and Homer1_SHANK (Figure 4D). Of those, only the dissociations of Cx36_NMDAR1 and Cx36_GluR2 were significant in HetB mice, and only the dissociation of Cx36_NMDAR1 was significant in HetC mice (Figure 4D-F, asterisks). Of note, HomB and HetC mice both showed significant dissociation of mGluR5 to a signaling effector (mGluR5_PIKE or mGluR5_Fyn).

**Figure 4.**
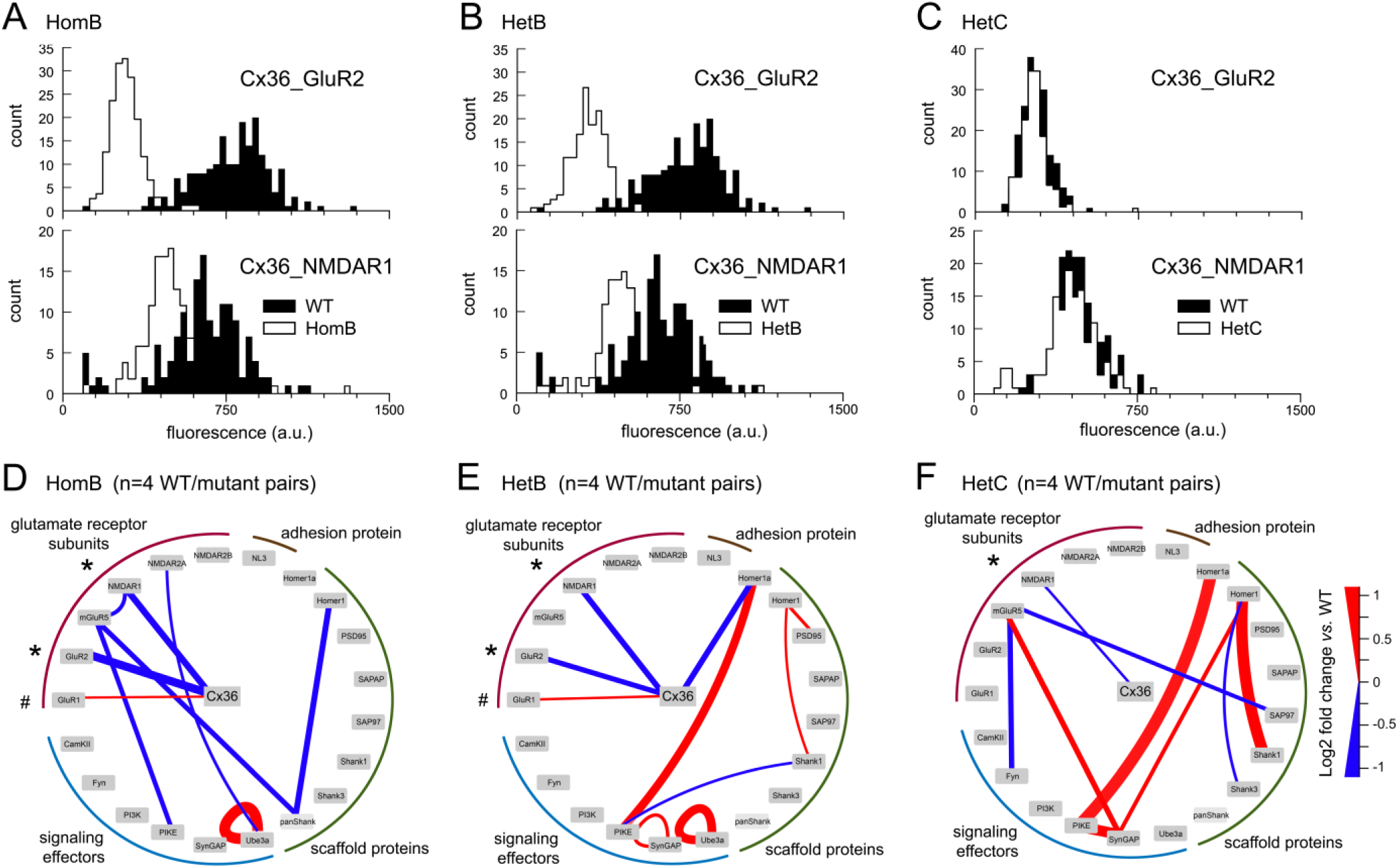
SHANK3 scaffolds Cx36 into the synaptic multiprotein network. (A-C) Example histograms of PiSCES Cx36_GluR2 and Cx36_NMDAR1 representative of N = 4 experiments. The left-shifted histograms of HomB and HetB mutants (white) indicate significantly (*p* < 0.05) less association of Cx36 to GluR2 and NMDAR1 compared to WT (black), with much weaker changes in a HetC mouse, assessed by KS test. (D-F) Node-edge diagrams showing the magnitude of all significantly changed PiSCES for each of the *Shank3* mutations (*p* < 0.05, assessed by ANC *∩* CNA). Edge thickness reflects the magnitude of change (red = increase; blue = decrease).

Protein dissociations due to *Shank3* mutations were accompanied by enhanced association among other proteins, indicating potential compensation. In HetB and HetC mice, the largest increases occurred for Homer1a_PIKE (HetB, 0.7 ± 0.3; HetC, 1.0 ± 0.1 log2-fold increase) and Homer1_Shank1 (each 0.4 ± 0.3 log2-fold increase). Both HomB and HetB mice demonstrated a small but significant increase in Cx36_GluR1 (HomB, 0.14 ± 0.03; HetB, 0.20 ± 0.04 log2-fold increase; Figure 4D,E, hash mark). Those findings indicated a restructuring of the synaptic multiprotein network by Homer1 and Shank1 due to *Shank3* mutations.

To summarize, Cx36 is structurally associated with the glutamatergic PSD *via* the PDZ binding domain of SHANK3 that links Cx36 to NMDAR1 and GluR2 subunits of ionotropic glutamate receptors. Dissociation of Cx36 from NMDAR1 occurs also with C-terminal truncation mutants of SHANK3 but at lower magnitude, and is the only dissociation common to both *Shank3* mutations. The results motivated studies of electrical synaptic transmission.

### *Shank3* mutations impair electrical synaptic transmission

Two key predictions from our results were that *Shank3* mutations should impair the modulation of electrical coupling by the NMDAR and should impair the synchrony of a network rhythm sustained by electrical synapses. We directly tested those predictions in three sets of electrophysiology studies of the *ex-vivo* IO using dual, whole-cell patch-clamp recordings of neighboring neuronal somata. The recordings were targeted to neurons no farther than 30 µm from one another, which we found was necessary to measure electrical coupling in the mouse IO. The first set of studies examined the effect of *Shank3* mutations on the baseline level of electrical coupling. Here, 74 pairs of IO neurons were studied from WT, HomB, HetB, and HetC mice.

In *a priori* control experiments, we examined the potential effect of *Shank3* mutations on three electrical properties of IO neurons that can influence coupling potentials and thereby confound comparisons of electrical coupling across genotypes. Because the junctional conductance (G_j_) mediated by electrical synapses is modulated by membrane conductances that can shunt current away from gap junctions into the extracellular space, we first assessed the input resistance (R_In_) of IO neurons under voltage clamp (−10 mV step). R_in_ was not statistically different among genotypes (WT, 135 ± 19; HetC, 82 ± 9; HetB, 174 ± 32; HomB, 143 ± 24 MΩ; *p* = 0.1, one-way ANOVA), indicating that the input conductance (G_in_ = R_in_) was unchanged and that differences in current shunting through the non-junctional membrane were unlikely to contribute to any change in electrical coupling in *Shank3* mutants. Second, because the membrane time constant of a neuron postsynaptic to electrical synapses determines the low-pass filtering of coupling potentials and, thus, electrical synapse throughput, we derived the membrane time constant from the slope of the linear relationship between membrane capacitance (C_m_) and G_in_. C_m_ was measured by the area of the capacitive transient to a −10 mV step and did not differ among genotypes (WT, 69 ± 9; HetC, 72 ± 8; HetB, 56 ± 9; HomB, 73 ± 7 pF; *p* = 0.6, one-way ANOVA) so that the time constant of IO neurons was unchanged by *Shank3* mutation (WT, 5.72; HetC, 5.08; HetB, 6.45; HomB, 6.48 ms). Similar C_m_ across genotypes indicated that the membrane surface area of IO neurons also was unchanged by *Shank3* mutations (WT, 6.94E-05 ± 9.27E-06 cm^2^). Third, we examined whether *Shank3* mutations affected the low-threshold calcium and hyperpolarization-activated cation currents (I_T_ and I_h_, respectively) that can influence coupling potentials due to their voltage activation near the resting membrane potential (Figure 5A). Across a range of hyperpolarizing voltage steps, peak I_h_ was measured as the difference between the instantaneous current after the capacitive transient at step onset and the steady-state current. Neither *Shank3* mutation altered I_h_ magnitude (post −50 mV step: WT, 54 ± 15; HetC, 68 ± 9; HetB, 36 ± 14; HomB, 79 ± 27 pA) or its voltage activation (Figure 5B). I_T_ was calculated from the peak of the tail current (I_tail_) of which 80% is due to I_T_ in mouse IO neurons (Figure 5A; Welsh et al., 2011). Neither *Shank3* mutation altered peak I_T_ (post −50 mV step: WT, 245 ± 30; HetC, 179 ± 34; HetB, 240 ± 108; HomB, 301 ± 67 pA) or its voltage activation (Figure 5B). Thus, *Shank3* mutations did not alter IO neurons’ intrinsic electrical properties in a manner that would influence coupling potentials mediated by electrical synapses.

**Figure 5.**
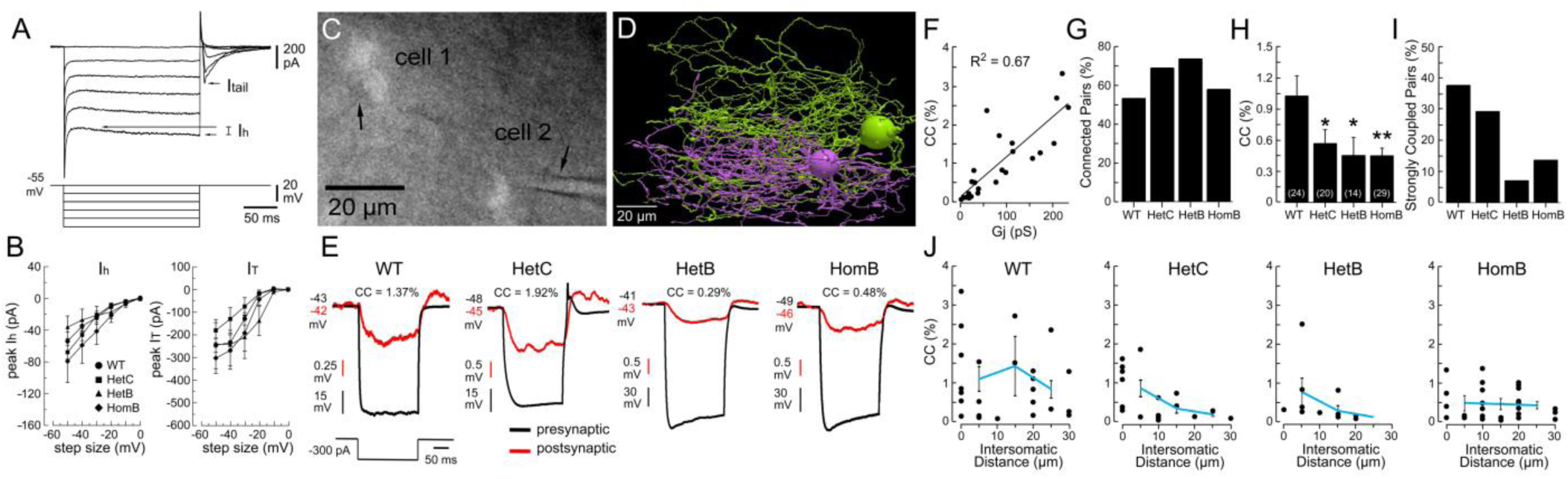
*Shank3* mutations impair electrical synaptic transmission. (A) Voltage clamp paradigm used to measure the magnitude and voltage-activation of I_h_ and I_T_ (5 hyperpolarizing steps, 10 mV increments from −55 mV holding potential). The low-threshold calcium current (I_T_) was calculated as 80% of the tail current (I_tail_) following step offset (Welsh et al., 2011). (B) I_h_ and I_T_ in IO neurons from *Shank3* mutants do not differ from WT in magnitude or voltage activation. Mixed-effects ANOVA did not detect a main effect of genotype for I_h_ magnitude (*p* = 0.5, *n* = 50 neurons) or I_T_ magnitude (*p* = 0.6, *n* = 28 neurons) or a genotype x voltage step interaction for I_h_ (*p* = 0.3) or I_T_ (*p* = 0.7). (C) Infrared differential-interference contrast image of the recording paradigm showing two patch electrodes (arrows) attached to the somata of two fluorescently labeled, neighboring IO neurons. (D) Three-dimensional reconstruction and surface rendering of the somata and dendritic arbors of a pair of electrically coupled neurons from the mouse IO. The image was obtained by dual intracellular biocytin and laser scanning confocal microscopy. Extensive dendritic overlap is a condition for dendrito-dendritic electrotonic coupling in IO neurons. (E) Examples of electrical coupling among neighboring IO neurons for all genotypes tested. Hyperpolarizing current was injected into a neuron presynaptic to electrical synapses (black) and a coupling potential was recorded in the postsynaptic neuron (red). (F) Linear relationship between G_j_ and CC for WT pairs of electrically coupled neurons. Each circle is an IO neuron postsynaptic to the electrical synapse. (G) Electrical coupling prevalence across genotypes. (H) Electrical coupling strength across genotypes (*n* = 87 neurons; mean CC ± SEM). One-way ANOVA indicated a main effect of genotype (*p* = 0.01). Dunnett’s *post hoc* tests indicated significant decreases in coupling strength from WT in all *Shank3* genotypes (HetC, *p* = 0.033; HetB, *p* = 0.015; HomB, *p* = 0.003). (I) Percentage of strongly coupled (> 1% CC) IO neurons for each genotype. (J) CC as a function of intersomatic distance for each genotype. Two-way ANOVA indicated a main effect of genotype to reduce CC (*p* = 0.004) but no main effect of intersomatic distance (*p* = 0.09) or genotype x intersomatic distance interaction (*p* = 0.6).

Electrical coupling was then directly tested by intracellular current injection into one of two simultaneously recorded IO neurons in order to measure coupling potentials mediated by electrical synapses (Figure 5C). Intracellular labeling and dendritic reconstruction of a pair of electrically coupled neurons in the mouse IO demonstrated the intertwined dendritic arbors that enable dendrito-dendritic electrical synapses (Sotelo et al., 1974, Hoge et al., 2011, Turecek et al., 2014; Figure 5D). Hyperpolarizing current injected into IO neurons presynaptic to electrical synapses produced a coupling potential in postsynaptic neurons whose magnitude was quantified by the ratio of the postsynaptic to presynaptic voltage responses (coupling coefficient, CC; Figure 5E). In WT mice, CC ranged between 0.065 and 3.25% and was linearly correlated to G_j_ (Figure 5F), indicating that CC scaled to electrical synapse strength under our conditions.

*Shank3* mutations decreased electrical coupling strength without reducing the prevalence of electrical coupling (Figure 5G,H). Mean CCs were: WT, 1.03 ± 0.19%; HetC, 0.57 ± 0.13%, HetB, 0.44 ± 0.17%, HomB, 0.44 ± 0.07%, corresponding to 57% weaker electrical coupling in Shank3B mutants (Figure 5H). The prevalence of strong electrical coupling (CC ≥ 1%) was reduced in HetB and HomB mice (7 and 14%, respectively) compared to WT (38%) and HetC (29%) mice (Figure 5I). WT mice exhibited uniform coupling strength over 0-30 µm intersomatic distance while HetC mice showed strong coupling only at somatic distance less than 10 µm (Figure 5J). Infrequent instances of strong electrical coupling in HetB and HomB mice were unrelated to somatic distance. *Shank3* mutations also affected the reciprocity of electrical coupling as 50% of WT pairs showed reciprocal coupling, while 35% of HetC, 43% of HetB, and 21% of HomB pairs showed reciprocal coupling.

### NMDAR strengthening of electrical coupling requires SHANK3 PDZ binding

In a second set of electrophysiology studies, we tested whether *Shank3* mutations impaired the plasticity of electrical coupling induced by NMDAR activation. Previous reports demonstrated that synaptic and pharmacological activation of NMDARs strengthened electrical coupling in the rat IO during an increase in G_in_, an increase in Ca^2+^ influx, and activation of CaMKII (Turecek et al., 2014; Welsh and Turecek, 2017). Here, we pharmacologically stimulated pairs of electrically coupled IO neurons with a weakly depolarizing concentration of NMDA (5 µM) to activate NMDARs equivalently across genotypes (depolarization by NMDA: WT, 4.6 ± 2.1; HetC, 0.2 ± 0.7; HetB, 1.3 ± 1.5; HomB, 1.3 ± 0.9 mV; *p* = 0.1, one-way ANOVA). A total of 69 electrically coupled IO neurons were studied.

On average, electrical coupling in WT and HetC mice showed a trend to become stronger during NMDAR activation (25 ± 27% and 108 ± 71% CC increase; 71 ± 61% and 120 ± 76% G_j_ increase, respectively). In contrast, IO neurons from HomB mice showed 18 ± 14% weaker electrical coupling during NMDAR activation, coincident with a 22 ± 16% decrease in G_j_ (Figure 6A,B). Figure 6C demonstrates the effects of NMDAR activation on electrical coupling in representative neuron pairs from WT and HomB mice. Of note, the heterogeneous responses of electrical coupling to the low concentration of NMDA precluded detecting statistically significant effects of genotype.

**Figure 6.**
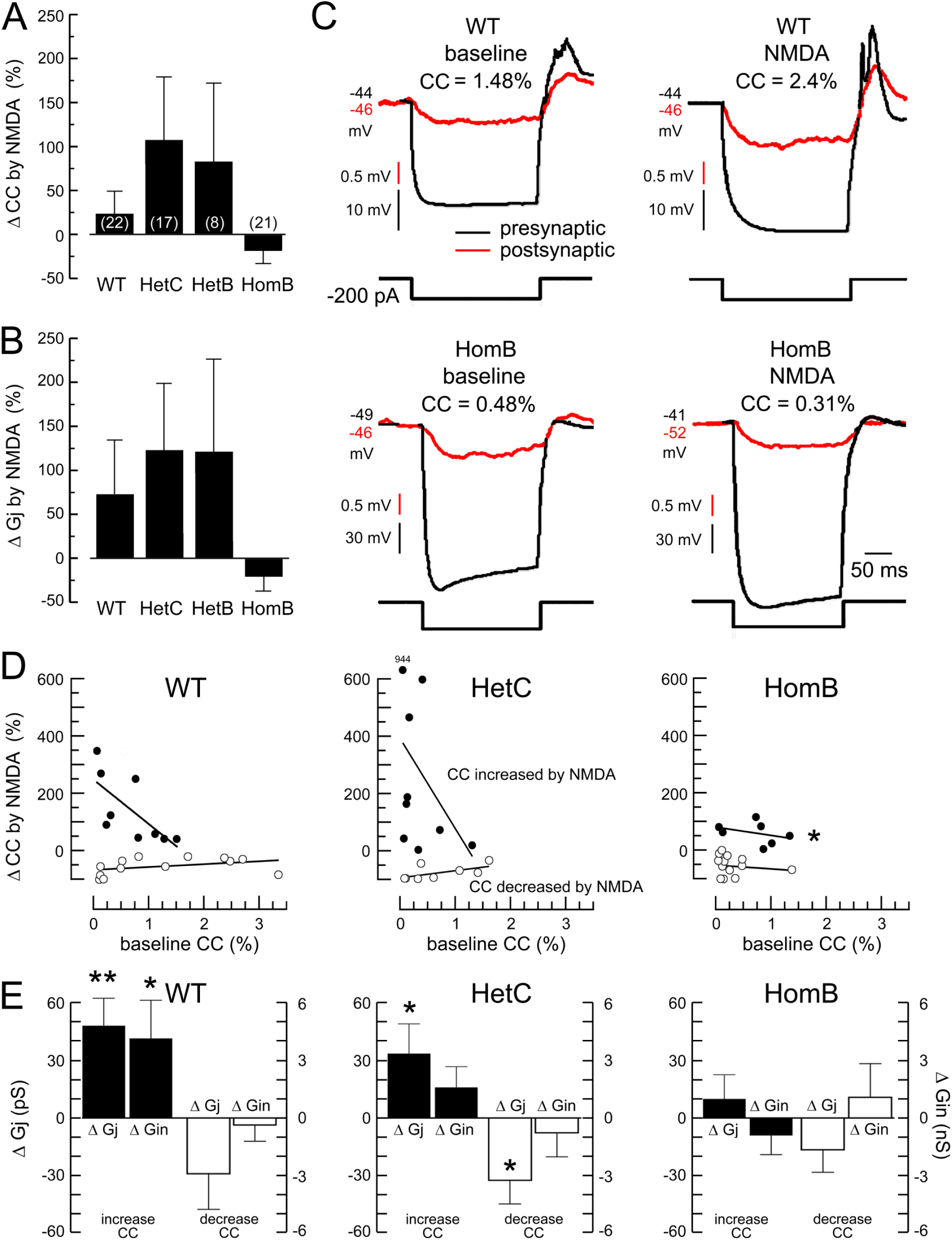
NMDAR strengthening of electrical coupling requires SHANK3 PDZ binding. (A,B) Mean percentage change in CC and G_j_ by 5 µM NMDA for WT and three *Shank3* mutant genotypes. Due to high variability, significant effects of genotype were not detected by one-way ANOVA (both *p* > 0.15). *N* = 68 electrically coupled IO neurons. (C) Representative recordings showing the strengthening and weakening of electrical coupling in IO neurons by NMDAR activation. (D) Strengthening of electrical coupling by NMDAR activation in WT and HetC mice was inversely related to electrical coupling strength at baseline, while HomB mice showed no significant strengthening of electrical coupling (filled circles). Statistically significant differences between HomB and WT mice (* *p* = 0.028) were identified by analysis of covariance (ANCOVA) but not between HetC and WT mice (*p* = 0.31). The weakening of electrical coupling in HetC and HomB mice did not differ from WT (*p* = 0.13, *p* = 0.40, respectively, by ANCOVA). Each circle shows an IO neuron that was electrically coupled at baseline. (E) Mean change in G_j_ and G_in_ during NMDAR activation for the neurons in D. HetC is unchanged from WT, but HomB shows no significant modulation of either conductance during electrical coupling strengthening. Asterisks indicate significant difference from zero by one-sample t-test (** *p* < 0.01, * *p* < 0.05).

To examine this further, we parsed the neurons from each genotype into two groups that showed strengthened or weakened electrical coupling during NMDAR activation. This was consistent with our previous report demonstrating bidirectional responses of electrical coupling to NMDAR activation in rat (Turecek et al., 2014) and a report of the mouse IO in which synaptic activation decreased coupling strength (Mathy et al., 2014). Parsing the neurons this way allowed us to examine the effect of *Shank3* mutations on both the strengthening and weakening of electrical coupling by NMDAR activation.

The Shank3B mutation blocked the strengthening of electrical coupling without affecting the weakening of electrical coupling induced by NMDAR activation. Conversely, the Shank3C mutation affected neither the strengthening nor weakening of electrical coupling. In WT mice, NMDAR-mediated strengthening of electrical coupling was inversely related to the baseline CC (Figure 6D, filled circles) and was positively related to increases in G_j_ and G_in_ (Figure. 6E), replicating previous report (Turecek et al., 2014). Those fundamental properties of strengthening plasticity of electrical coupling were unchanged in magnitude and form in HetC mice. Yet, HomB mice showed neither significant strengthening of electrical coupling nor an increase in G_j_ with NMDAR activation. Weakened electrical coupling by NMDAR activation was similar across genotypes in the overall magnitude of the decrease, its independence from baseline coupling strength, and relation to a reduction in G_j_ (Figure 6D,E)

As predicted from the proteomics, the Shank3B mutation had the greatest negative impact on the modulation of electrical coupling by NMDAR activation, consistent with its effect to produce the strongest dissociation of Cx36_NMDAR1. The experiment provided the unexpected insight that only strengthening of electrical coupling was impaired by *Shank3* mutations, indicating that NMDAR-induced weakening of electrical coupling does not depend upon the Cx36-PSD interactions revealed by the proteomics.

### *Shank3* mutations disrupt brain rhythm synchrony

The third set of electrophysiology studies tested whether *Shank3* mutations that disrupt the strength and plasticity of electrical synaptic transmission also disrupt the synchrony of a electrical brain rhythm. Electrical synapses among IO neurons synchronize oscillations in membrane potential that are below the threshold for action potentials (Llinás and Yarom, 1986; Long et al., 2002; Figure 7A). Such subthreshold oscillations (STOs) produce recurring epochs of increased spike probability that drive coherent activity in the cerebellum (Welsh et al., 1995; Marshall et al., 2007). Biophysical modeling indicated that STOs are synchronized by a recurring depolarizing current that is conducted through the IO network *via* electrical synapses (Manor et al., 1997). Electrophysiology affirmed that blocking electrical coupling prevents STO synchrony (Long et al., 2002) and reduces STO amplitude (Placantonakis et al., 2006) while strengthening electrical coupling by NMDAR activation increases STO synchrony and amplitude (Turecek et al., 2014; Welsh and Turecek, 2017). STOs were recorded in 46 pairs of IO neurons.

**Figure 7.**
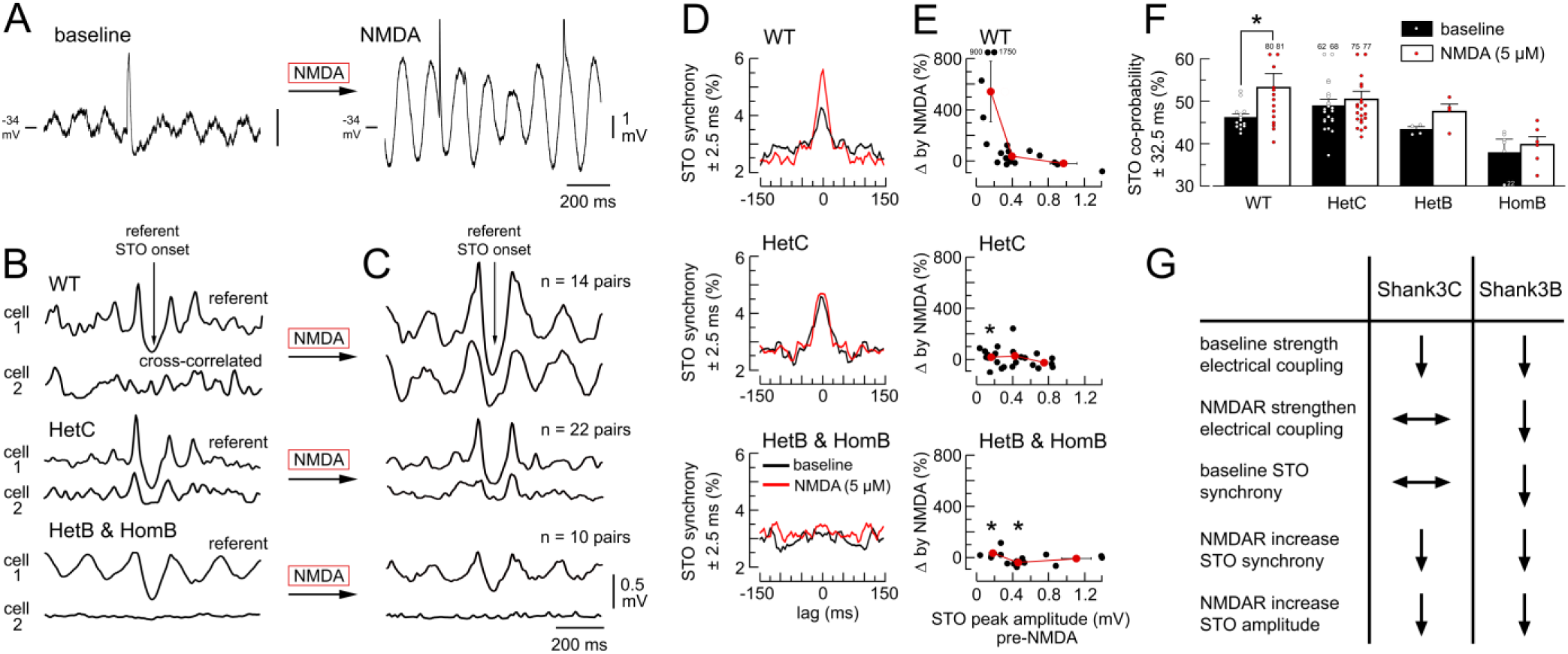
*Shank3* mutations disrupt brain rhythm synchrony. (A) Membrane potential of a mouse IO neuron shows STOs at baseline and the effect of NMDAR activation to enhance STO amplitude. Note the action potentials (truncated for clarity) riding on the depolarized phases of the STO. (B,C) Average membrane potential of two simultaneously recorded IO neurons within 30 µm of one another for WT mice and *Shank3* mutants. For each genotype, averaging is synced to STO onset of cell 1 (downward arrow), producing an auto-correlated trace for cell 1 and a cross-correlated trace for cell 2. NMDAR activation enhances the synchrony, amplitude, and regularity of simultaneously recorded STOs in WT mice but not in Shank3C or Shank3B mutants. Robust STO correlation was not observed in the average traces of HomB mice before or during NMDAR activation. Recordings were obtained from 46 pairs of IO neurons. (D) Mean cross-correlograms of STO onsets show STO onset synchrony (time 0) before (black) and during (red) NMDAR activation for WT and *Shank3* mutants. Each plot shows the relative timing of STO onsets (± 150 ms) for the traces comprising the averages in B and C. (E) Percentage change in STO amplitude by NMDAR activation as a function of baseline. Red curves show mean ± SEM in tertile averages. Asterisks indicate significant difference compared to WT (*p* < 0.05 by one-way ANOVA and Dunnett’s *post hoc* test). (F) STO co-occurrence (± 32.5 ms) before and during NMDAR activation. NMDAR activation enhances STO co-occurrence in WT mice only. Each dot is a single IO neuron. Two-way ANOVA indicated main effect of genotype (*p* = 0.008) and NMDA (*p* = 0.018). *Post-hoc* tests within group indicated a significant effect of NMDA only in WT (* *p* = 0.037). (G) Summary of *Shank3* mutation effects on electrical coupling and STO synchrony in the IO.

The Shank3B mutation, but not the Shank3C mutation, reduced the baseline level of STO synchrony as seen in cross-correlated membrane potentials (Figure 7B) and STO onset cross-correlograms (Figure 7D, black). The absence of STO synchrony due to the Shank3B mutation agreed with the 57% reduction in electrical coupling strength at baseline. Of note, STO synchrony of HetC mice was equivalent to WT at baseline, indicating that the preservation of strong electrical coupling was sufficient to support STO synchrony (Figure 5I).

Both *Shank3* mutations prevented NMDAR activation from enhancing STO synchrony (Figure 7C-E). STOs of WT mice became more synchronized during NMDAR activation but neither *Shank3* mutation showed this effect (Figure 7C-E). Thus, for Shank3B mutants, STOs remained as unsynchronized during NMDAR activation as they were at baseline. For Shank3C mutants, STO synchrony remained at the baseline level during NMDAR activation. NMDAR activation increased STO amplitude in WT mice, primarily by enhancing STOs on the low end of the peak amplitude range (< 0.3 mV) from 0.2 ± 0.03 to 1.0 ± 0.5 mV (Figure 7F). In contrast, NMDAR activation did not enhance STO amplitude in either HetC or HetB/HomB mice.

To summarize, the findings demonstrated that *Shank3* mutations blocked STO synchronization and the enhancement of STOs by NMDAR activation, a property of the IO network that is underlain by the strengthening of electrical synaptic transmission (Turecek et al., 2014; Welsh and Turecek, 2017). The Shank3B mutation produced the most deleterious phenotype by showing no STO synchrony. The Shank3C mutation produced a less severe phenotype in which STO synchrony was normal at baseline but could not be upregulated by NMDAR activation. Figure 7G summarizes the electrophysiological effects of *Shank3* mutations on electrical synaptic transmission and the synchrony of an electrical rhythm within the IO network.

## DISCUSSION

The experiments demonstrated that Cx36, the protein that comprises neuronal electrical synapses, is structurally associated with the glutamatergic PSD in mouse brain by the scaffold protein SHANK3. That two different mutations of *Shank3* would reduce the strength of electrical synaptic transmission was not predicted prior to the proteomic experiments performed here. The finding that Cx36 is integrated into the synaptic multiprotein network by SHANK3 advances our understanding of synaptic transmission (Pereda, 2014; Nagy et al., 2018). Additionally, our experiments demonstrate a mechanism by which disruptions of the glutamatergic synaptic multiprotein network by *Shank3* mutations can impair an intrinsic electrical rhythm in the brain. Because the ability of the brain to generate electrical rhythms is fundamental to cognition and is supported by the functioning of electrical synapses (Buhl et al., 2003; Coulon and Landisman, 2017; Llinás, 2014; Pernelle et al., 2018; Traub et al., 2020; Welsh et al., 2005; Welsh and Turecek, 2017), our findings help establish the proteomic/electrophysiological link between *Shank3* mutations and ID.

### Cx36 complexes with NMDAR1 and GluR2 in an “extended PSD”

A structural relationship between neuronal electrical synapses and the glutamatergic PSD was first inferred based on the anatomy and physiology of a specialized junction made by the 8^th^ cranial nerve upon the reticulospinal Mauthner cell in goldfish (Faber et al., 1980; Pereda et al., 1994, 2003; Robertson et al., 1963). That unique junction was termed a “mixed synapse” because it contains both a gap junction plaque and a glutamatergic PSD within the postsynaptic membrane of the axon terminal bouton, an arrangement found sporadically in the mammalian brain (Nagy et al., 2018). Gap junctions similarly positioned relative to glutamatergic PSDs were later discovered in the rat IO (Hoge et al., 2011; Turecek et al., 2014). The subcellular anatomy of the two structures differed, however, as the gap junction in the teleost mixed synapse is axo-somatic and within the post-synaptic membrane of the terminal bouton while gap junctions near PSDs in the mammalian IO are dendrito-dendritic and outside the terminal bouton. Thus, as an anatomical motif, the PSD-gap junction assembly is implemented in different ways to accomplish different functions: in the mixed synapse, modulating the gap junction regulates the throughput of a single axon; in the mammalian IO, modulating the gap junction regulates communication between adjacent neurons. It is important that the PSD-gap junction assembly in the IO provides a highly efficient mechanism by which chemical synaptic transmission can modulate ongoing network activity that is unconstrained by the episodic and relatively slow speed of axonal conduction (Llinás, 2014).

QMI revealed a set of PiSCES that helps explain the proximity of neuronal gap junctions to the glutamatergic PSD and which justifies considering electrical synapses as components of an “extended PSD.” The extended PSD is defined minimally by strong associations between Cx36 and the ionotropic glutamate receptor subunits GluR2 and NMDAR1. The importance of SHANK3 for scaffolding Cx36 to GluR2 is consistent with a SAYV PDZ interaction motif on the Cx36 C-terminus (Li et al., 2004, 2012) and the binding of SHANK3 to GluR2 (Ponna et al., 2018). Based on known interactions, SHANK3 may complex with NMDAR1 by binding to the scaffold protein SAPAP which in turn binds to the PSD-95_NMDAR complex (Monteiro and Feng, 2017). Although not examined in our study, the scaffold protein ZO-1 interacts with Cx36 and can interact with SHANK3, providing another way by which Cx36 may be integrated into the synaptic multiprotein complex (Collins 2006; Li et al., 2004). That the distance between neuronal gap junctions and glutamatergic PSDs is on the order of 100-200 nm (Hoge et al., 2011; Pereda et al., 2003) indicates that multiple dimers of SHANK3 interacting partners, perhaps in combination with Homer tetramers (Hayashi et al., 2009), mediate the Cx36 associations described here, given the 60 nm length of SHANK3 (Kursula, 2019). Of note was that the reduced association of Cx36 to NMDAR1 and GluR2 in Shank3B mutant mice was accompanied by the strengthening of an association between Cx36 and GluR1, indicating that there can be multiple configurations of the multi-protein scaffold that integrates Cx36 to the PSD.

### Graded effects of *Shank3* mutations on IO network synchronization

Heterogeneity in the synaptic protein interactome following two *Shank3* mutations indicated that different rearrangements in the associations among glutamate receptors, signaling effectors, and electrical synapses will impart diversity to the consequent behavioral and molecular deficits (Monteiro and Feng, 2017). Only the reduction in the association between Cx36 and NMDAR1 was common to the Shank3B and Shank3C mutations. That result reinforced our main conclusion that Cx36 is integrated into the PSD multiprotein network and that electrical synaptic transmission is supported by SHANK3.

Although dissociation Cx36 from NMDAR1 was common to two *Shank3* mutations, differences in magnitude of that dissociation related to different changes in the properties of electrical coupling. Changes in electrical coupling following *Shank3* mutations suggest different numbers of gap junctions within the dendritic arbor, different numbers of intercellular channels within gap junctions, and/or a change in the single channel conductance of electrical synapses. The former two possibilities could follow from impairments in the stabilization of gap junctions and their intercellular channels in the plasmalemma membrane. The latter possibility would implicate a constitutive action of NMDARs and intracellular signaling to support the baseline level of electrical coupling. Phosphorylation of Cx36 by CaMKII at a consensus site near the Cx36 C-terminus increased the single channel conductance of neuronal electrical synapses (Del Corsso et al., 2012). The sensitivity of Cx36 to local domains of Ca^2+^ influx and activated CaMKII underlies activity-dependent strengthening of electrical synapses in IO neurons (Turecek et al., 2014; Welsh and Turecek, 2017). Recent work demonstrated that phosphorylation of the CaMKII consensus site within Cx36 also increases PDZ binding (Tetenborg et al., 2020). The combined evidence implicates a feed-forward process in which CaMKII activation simultaneously increases electrical synapse strength and stabilizes gap junction intercellular channels within the extended PSD *via* interactions involving Cx36, SHANK3, and NMDAR1.

The *Shank3* mutations we studied impaired STO synchronization to degrees that scaled with the reductions in electrical coupling. The Shank3B mutation produced the greatest dissociation of Cx36 from NMDAR1, the greatest impairment of electrical coupling at baseline, and prevented NMDAR activation from strengthening electrical coupling, while desynchronizing STOs and blocking synchronization by NMDAR activation. The Shank3C mutation produced a lesser dissociation of Cx36 from NMDAR1 and a lesser reduction in electrical coupling, but did not block NMDAR activation from strengthening electrical coupling. Yet, the enhancement of electrical coupling by NMDAR activation in Shank3C mutants was insufficient to enhance network synchrony, perhaps due to lower mean coupling strength within the population or other changes within the synaptic interactome whose contribution is currently unclear. Indeed, the upregulated associations of signaling effectors with each other, with other scaffold proteins, and with mGluR5 are likely to contribute to physiological phenotypes after *Shank3* mutation. To summarize, the effects of *Shank3* mutations on IO network synchronization depended upon the site of mutation and the signature disruptions of the synaptic protein interactome unique to each mutation.

### Implications for intellectual function

Our findings indicate that effects of *Shank3* mutations on electrical synaptic transmission may contribute to core symptoms in subpopulations of individuals with ASD and schizophrenia. This will be manifested as desynchronized 4-10-Hz olivo-cerebellar oscillations whose coherence helps encode the cerebellar contribution to motor, learning, and cognitive function (Schmahmann, 2019; Tsutsumi et al., 2020; Welsh et al., 2005; Welsh and Turecek, 2017; Wu et al., 2011) and whose biophysical origin was studied here directly. The results are likely to apply to effects of *Shank3* mutations on the generation of coherent oscillations elsewhere in the brain that contribute to intellectual functioning. For instance, 30-70 Hz gamma-band oscillations in the cerebral cortex contribute to cognition and arise from the modulation of electrical coupling among inhibitory interneurons (Traub et al., 2001) and the fine balance of glutamatergic neurotransmission among deep and superficial layers of neocortex (Ainsworth et al., 2011), both of which will be affected by *Shank3* mutations. Consistently, the severity of symptom expression in ASD and schizophrenia scaled to the degree of impairment in the generation of gamma band oscillation (Edgar et al., 2015; Grent-’t-Jong et al., 2018). Furthermore, ASD and schizophrenia are commonly associated with disturbances in the sleep-wake cycle (Ashton and Jagannath, 2020; Ballester et al., 2020), of which transitions in sleep states may require modulation of electrical synapses in the reticular thalamus by chemical synaptic transmission (Coulon and Landisman, 2017). The foregoing indicates that agents which upregulate electrical synaptic transmission (Urbano et al., 2007; Beck et al., 2008) may benefit individuals with syndromes due to *Shank3* mutations.

## AUTHOR CONTRIBUTIONS

J.L., Z.Z., H.E.S. performed experiments. J.L., H.E.S., S.E.P.S., and J.P.W. designed experiments and wrote the paper.

## ACKNOWLEDGMENTS

Supported by NIH Grants R01 NS31224-24 and R01 MH113545. We thank Dr. S. Wang for Shank3C mice.

## STAR METHODS

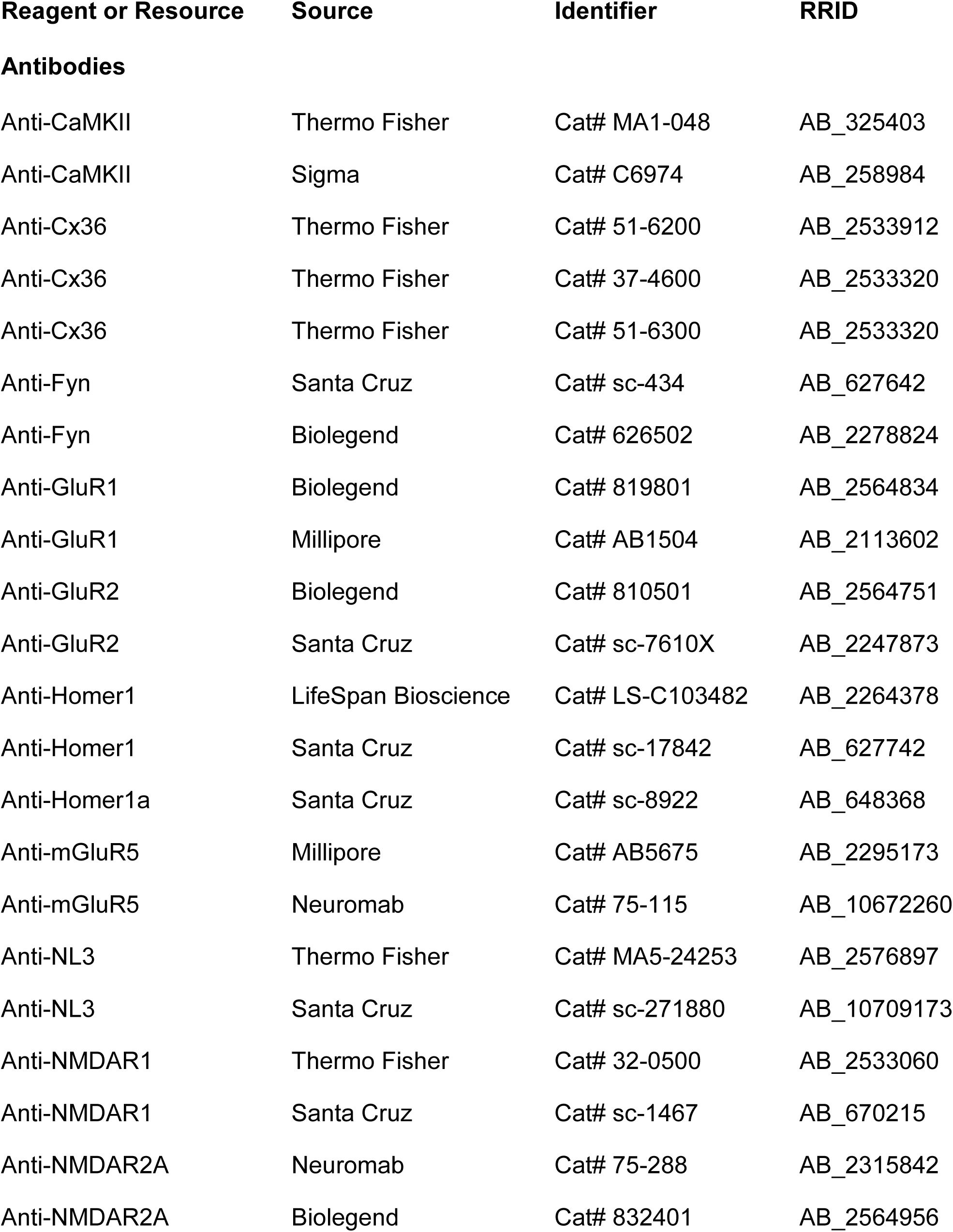
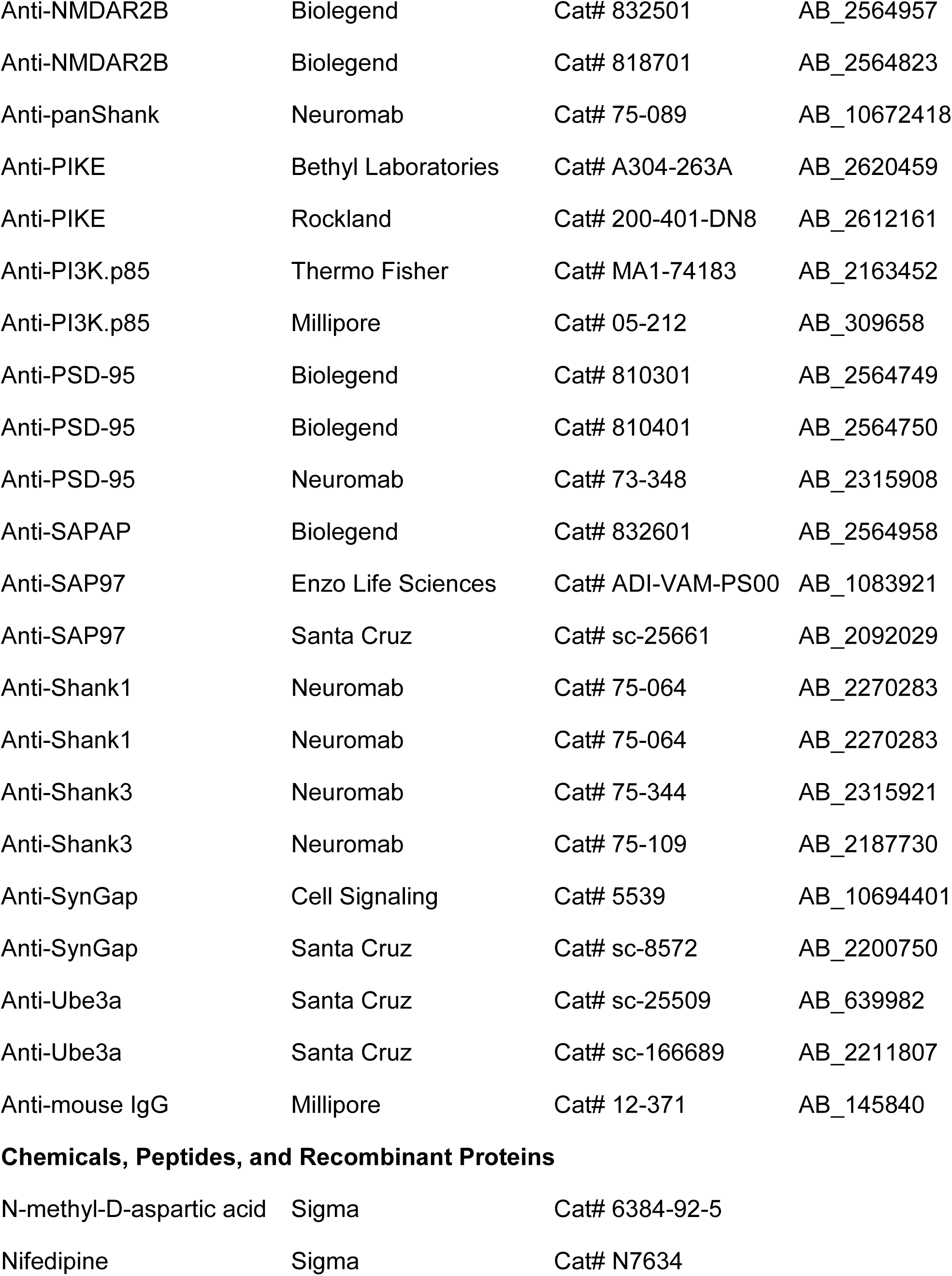
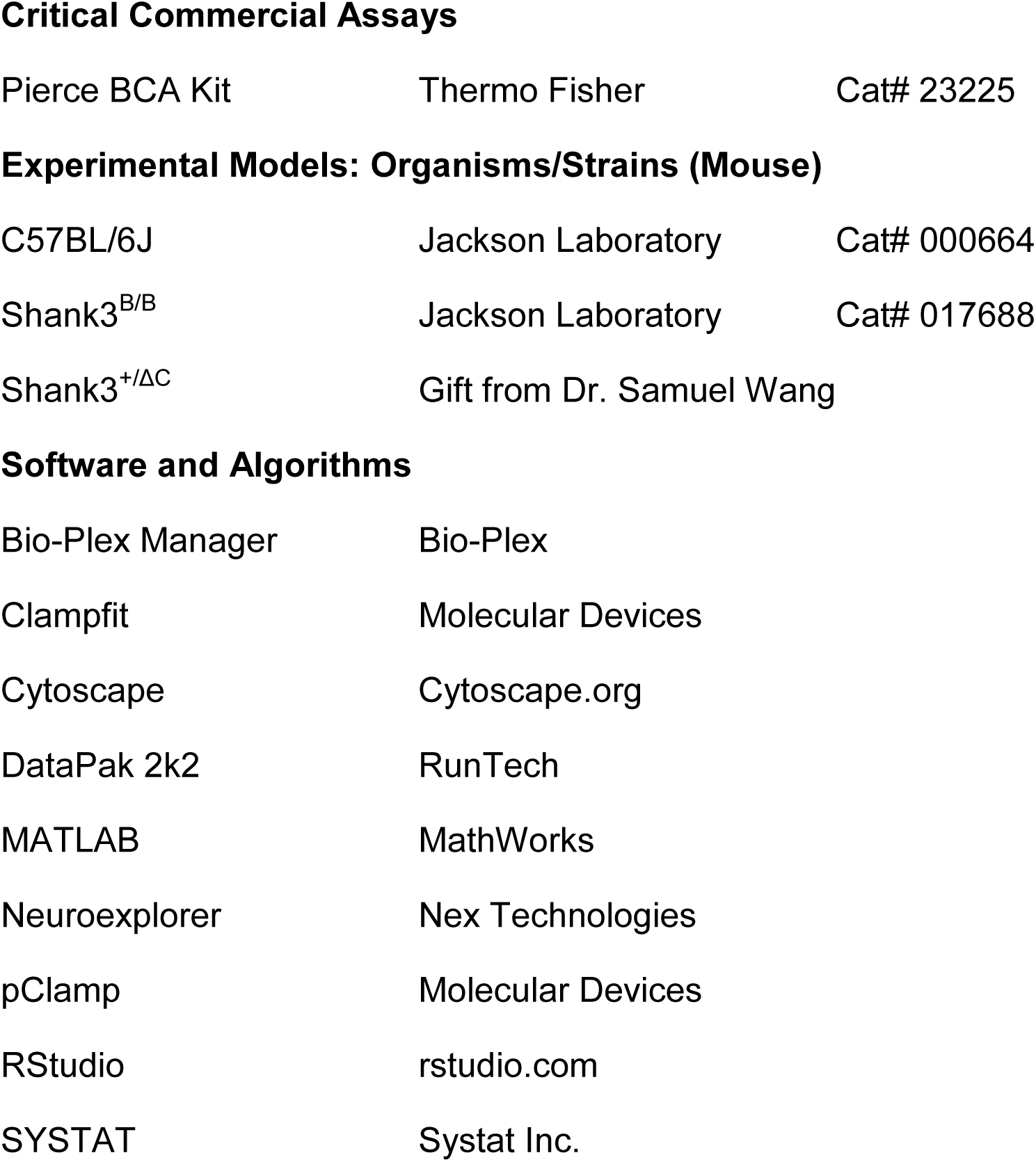

| **Reagent or Resource** | **Source** | **Identifier** | **RRID** |
| --- | --- | --- | --- |
| **Antibodies** |  |  |  |
| Anti-CaMKII | Thermo Fisher | Cat# MA1-048 | AB_325403 |
| Anti-CaMKII | Sigma | Cat# C6974 | AB_258984 |
| Anti-Cx36 | Thermo Fisher | Cat# 51-6200 | AB_2533912 |
| Anti-Cx36 | Thermo Fisher | Cat# 37-4600 | AB_2533320 |
| Anti-Cx36 | Thermo Fisher | Cat# 51-6300 | AB_2533320 |
| Anti-Fyn | Santa Cruz | Cat# sc-434 | AB_627642 |
| Anti-Fyn | Biolegend | Cat# 626502 | AB_2278824 |
| Anti-GluR1 | Biolegend | Cat# 819801 | AB_2564834 |
| Anti-GluR1 | Millipore | Cat# AB1504 | AB_2113602 |
| Anti-GluR2 | Biolegend | Cat# 810501 | AB_2564751 |
| Anti-GluR2 | Santa Cruz | Cat# sc-7610X | AB_2247873 |
| Anti-Homer1 | LifeSpan Bioscience | Cat# LS-C103482 | AB_2264378 |
| Anti-Homer1 | Santa Cruz | Cat# sc-17842 | AB_627742 |
| Anti-Homer1a | Santa Cruz | Cat# sc-8922 | AB_648368 |
| Anti-mGluR5 | Millipore | Cat# AB5675 | AB_2295173 |
| Anti-mGluR5 | Neuromab | Cat# 75-115 | AB_10672260 |
| Anti-NL3 | Thermo Fisher | Cat# MA5-24253 | AB_2576897 |
| Anti-NL3 | Santa Cruz | Cat# sc-271880 | AB_10709173 |
| Anti-NMDAR1 | Thermo Fisher | Cat# 32-0500 | AB_2533060 |
| Anti-NMDAR1 | Santa Cruz | Cat# sc-1467 | AB_670215 |
| Anti-NMDAR2A | Neuromab | Cat# 75-288 | AB_2315842 |
| Anti-NMDAR2A | Biolegend | Cat# 832401 | AB_2564956 |
| Anti-NMDAR2B | Biolegend | Cat# 832501 | AB_2564957 |
| Anti-NMDAR2B | Biolegend | Cat# 818701 | AB_2564823 |
| Anti-panShank | Neuromab | Cat# 75-089 | AB_10672418 |
| Anti-PIKE | Bethyl Laboratories | Cat# A304-263A | AB_2620459 |
| Anti-PIKE | Rockland | Cat# 200-401-DN8 | AB_2612161 |
| Anti-PI3K.p85 | Thermo Fisher | Cat# MA1-74183 | AB_2163452 |
| Anti-PI3K.p85 | Millipore | Cat# 05-212 | AB_309658 |
| Anti-PSD-95 | Biolegend | Cat# 810301 | AB_2564749 |
| Anti-PSD-95 | Biolegend | Cat# 810401 | AB_2564750 |
| Anti-PSD-95 | Neuromab | Cat# 73-348 | AB_2315908 |
| Anti-SAPAP | Biolegend | Cat# 832601 | AB_2564958 |
| Anti-SAP97 | Enzo Life Sciences | Cat# ADI-VAM-PS00 | AB_1083921 |
| Anti-SAP97 | Santa Cruz | Cat# sc-25661 | AB_2092029 |
| Anti-Shank1 | Neuromab | Cat# 75-064 | AB_2270283 |
| Anti-Shank1 | Neuromab | Cat# 75-064 | AB_2270283 |
| Anti-Shank3 | Neuromab | Cat# 75-344 | AB_2315921 |
| Anti-Shank3 | Neuromab | Cat# 75-109 | AB_2187730 |
| Anti-SynGap | Cell Signaling | Cat# 5539 | AB_10694401 |
| Anti-SynGap | Santa Cruz | Cat# sc-8572 | AB_2200750 |
| Anti-Ube3a | Santa Cruz | Cat# sc-25509 | AB_639982 |
| Anti-Ube3a | Santa Cruz | Cat# sc-166689 | AB_2211807 |
| Anti-mouse IgG | Millipore | Cat# 12-371 | AB_145840 |

**Chemicals, Peptides, and Recombinant Proteins**
|  |  |  |
| --- | --- | --- |
| N-methyl-D-aspartic acid | Sigma | Cat# 6384-92-5 |
| Nifedipine | Sigma | Cat# N7634 |

Critical Commercial Assays
|  |  |  |
| --- | --- | --- |
| Pierce BCA Kit | Thermo Fisher | Cat# 23225 |

Experimental Models: Organisms/Strains (Mouse)
|  |  |  |
| --- | --- | --- |
| C57BL/6J | Jackson Laboratory | Cat# 000664 |
| Shank3<sup>B/B</sup> | Jackson Laboratory | Cat# 017688 |

|  |  |
| --- | --- |
| Shank3<sup>+/ΔC</sup> | Gift from Dr. Samuel Wang |

Software and Algorithms
|  |  |
| --- | --- |
| Bio-Plex Manager | Bio-Plex |
| Clampfit | Molecular Devices |
| Cytoscape | Cytoscape.org |
| DataPak 2k2 | RunTech |
| MATLAB | MathWorks |
| Neuroexplorer | Nex Technologies |
| pClamp | Molecular Devices |
| RStudio | rstudio.com |
| SYSTAT | Systat Inc. |

## RESOURCE AVAILABILITY

### Lead Contact

Further information and requests for resources and reagents should be directed to and will be fulfilled by the Lead Contact, J. Welsh.

### Materials Availability

This study did not generate new unique reagents.

### Data and Code Availability

The data and custom code that support the findings from this study are available from the Lead Contact upon request.

## EXPERIMENTAL MODEL AND SUBJECT DETAILS

### Animals

Procedures were performed in accordance with the U.S. NIH guidelines for the care and use of laboratory animals and were approved by the Seattle Children’s Animal Care and Use Committee. Mice both male and female were used (age P28-P60). C57BL/6J founder mice were purchased from The Jackson Laboratory. Shank3C founders were obtained from Dr. Samuel Wang. Shank3B founders were obtained from The Jackson Laboratory. The genotype of every experimental mouse was verified by PCR from skin punch.

## METHOD DETAILS

### PSD preparation

The IO of P60 mice was dissected from medulla oblongata and homogenized by a glass teflon homogenizer in sucrose buffer (0.32 M sucrose, 4 mM HEPES, pH 7.4, protease/phosphatase inhibitors, Sigma, P8340/P5726). Whole cell lysate was spun (1000 *g*, 10 min, 4⁰ C), the P1 pellet was homogenized with 200 µl membrane solubilizing buffer (mM): HEPES 50, EGTA 2, Na3VO4 5, NaF 30, 1% Triton X-100, protease/phosphatase inhibitors). The S1 was spun (15,000 *g,* 15 min, 4⁰ C), the supernatant (S2) saved, and the P2 pellet was lysed in 300 µl membrane solubilizing buffer and spun (150,000 *g*, 40 min, 4⁰ C) before being placed in 50 µl of PSD solubilizing buffer (mM): (Tris 50, NaF 30, Na3VO4 5, ZnCl2 0.02, Na-deoxycholate 1%, protease/phosphatase inhibitors).

### Co-IP and Western immunoblot

Mouse IO was homogenized in lysis buffer (mM): NaCl 150, Tris 50, 1% NP-40, NaF 1, Na3VO4 1, protease/phosphatase inhibitors, pH 7.4), followed by lysis buffer incubation (15 min on ice) and high-speed centrifugation (13,000 *g*, 15 min, 4⁰ C). Supernatant protein concentration was determined (Pierce BCA kit 23225). Protein (200 µg) was precleared with protein G magnetic beads (New England Biolabs) and incubated overnight (4° C) with 5 µg of Cx36 (Thermofisher, 37-4600, RRID: AB_2533320) or NMDAR1 (Thermofisher, 32-0500, RRID: AB_2533060) IP antibody, or normal mouse IgG control (Millipore, 12-371, RRID: AB_145840), followed by protein G bead incubation (2.5 h, 25 µl/sample). After TBST washing (5x, 0.2 M Tris, 1.0 M NaCl, 0.1% Tween 20, 7.4 pH), samples were boiled in 4x Laemmli sample buffer and subjected to SDS-PAGE electrophoresis followed by western immunoblotting. Gels were transferred onto PVDF (Millipore) membranes, blocked in 5% milk in TBST (60 min), and primary antibodies applied overnight in blocking medium (Cx36, Thermofisher 51-6300, RRID: AB_2533913, 1:1000; NMDAR1, Thermofisher 32-0500, RRID: AB_2533060, 1:1000; PSD-95, Neuromab 73-348, RRID: AB_2315908, 1:2000). Blots were imaged (Protein Simple; Pierce Femto reagents) after washing and probing with species-specific HRP-conjugated antibodies.

### QMI

A 19 × 22 array of IP and probe antibodies was used (Supplemental Information) to measure 418 IP-probe combinations (Smith et al., 2016; Lautz et al., 2018, 2019). A master mix containing equal numbers of IP antibody-coupled Luminex beads was prepared and distributed into post-nuclear cell lysates in duplicate. Protein complexes were IP’d overnight from lysates containing equal amounts of protein, washed (2x Fly-P buffer), and distributed into a 96-well plate at twice the number of probes. Biotinylated probe antibodies were added and incubated with gentle agitation (1 h, 600 rpm). Microbeads and captured complexes were washed (3x Fly-P buffer, Bio-Plex Pro II magnetic plate washer), incubated with streptavidin-PE (30 min), washed (3x), and resuspended (125 μl Fly-P buffer). Fluorescence was measured by a custom Bio-Plex 200 (Bio-Plex Manager software, v. 6.1). Procedures were performed at 4° C or on ice. Data were processed to: 1) eliminate doublets on the basis of the doublet discriminator intensity (> 5000 and < 25 000 arbitrary units); 2) identify specific bead classes within the bead regions used; and 3) pair individual bead PE fluorescence measurements with their corresponding bead regions. The method generated a fluorescence intensity distribution for each paired measurement.

### Brain-stem slice electrophysiological recordings

Mice (P28-P56, both sexes) were anesthetized with isoflurane, decapitated, and brains placed into ice-cold solution containing (mM): sucrose 219, KCl 5, NaH_2_PO_4_ 1.25, NaHCO_3_ 26, dextrose 10, CaCl_2_ 0.5, MgSO_4_, oxygenated with 95% O_2_/5% CO_2_ (pH 7.4; 300-315 mOsm). Parasagittal brainstem slices (Leica VT1000 S) were transferred to recording ACSF containing (mM): NaCl 126, KCl 5, NaH_2_PO_4_ 1.25, NaHCO_3_ 26, dextrose 10, CaCl_2_ 2, MgSO_4_ 2, 95% O_2_/5% CO_2_ (pH 7.4, 300-315 mOsm), brought to 35⁰ C over 15 min, and maintained at room temperature for ≥ 1.5 h before recording (34⁰ C; 2-3 ml/min perfusion; Olympus BX-51WI microscope; Molecular Devices Multiclamp 700B amplifier; Digidata 1440 10-kHz digitization). Patch pipettes (6-8 MΩ) were filled with solution (mM): K gluconate 130, EGTA 5, HEPES 10, KCl 5, CaCl_2_ 0.5, MgSO_4_ 2, Na_2_ ATP 4, Na_2_ phosphocreatine 5, Na_3_ GTP 0.3 (pH 7.4, 289 mOsm). Paired whole-cell somatic recordings were obtained from IO neurons having somata no farther than 30 µm apart using two micromanipulators (Luigs and Neumann, B3ES).

Electrical coupling was quantified by injecting hyperpolarizing current (−50 to −500 pA, 200 ms, 0.125 Hz) into one of 2 neurons during dual, whole-cell patch recording, measuring both voltage responses, and calculating their ratio to derive the CC (Bennett, 1966; Devor and Yarom, 2002). CCs were calculated from averages (50-200 trials) and were measured from the average voltage 100 ms before current offset to minimize contribution of capacitive coupling. For CC measurements, NiCl_2_ or Nifedipine (50 µM each) were added to the recording solution to prevent STOs when needed (Turecek et al., 2014). G_j_ was calculated as: G_j_ = 1/[[(R_in_ cell 1)*(R_in_ cell 2)–(transfer resistance)^2^]/transfer resistance] (Bennett, 1966). Transfer resistance was defined as the voltage response in cell 2 when current was injected into cell 1 divided by the amplitude of the current step. R_in_ was calculated from a −10 mV voltage step.

Under voltage clamp, I_h_ and I_tail_ were elicited using a range of 200-ms hyperpolarizing voltage commands from a holding potential of −55 mV to a maximum of −105 mV in 10-mV increments before return to −55 mV. Peak I_h_ was measured as the difference between the instantaneous current, immediately after the capacitive transient following voltage command onset and the steady-state current in the 20 ms before the return step. Peak I_tail_ was measured as the maximum inward current occurring within 80 ms after offset of the voltage command using the current value before the voltage pulse as the baseline. Peak I_T_ was calculated from the peak of I_tail_ of which 80% is due to I_T_ in mouse IO neurons (Figure 5A; Welsh et al., 2011). Input resistance and membrane capacitance were assessed with the −10 mV step.

STOs were measured from 12-Hz low-pass filtered voltage records and were identified after differentiation to identify dV/dT peak events that corresponded to the beginning of the depolarizing phase of each STO. The events were time-stamped for auto- and cross-correlation analyses (DataPak 2k2; Neuroexplorer v. 5.131). STO amplitudes were measured from average records for each IO neuron.

## QUANTIFICATION AND STATISTICAL ANALYSIS

### Adaptive non-parametric analysis with weighted cutoff (ANC) and correlation network analysis (CNA)

This analysis pipeline was developed for QMI data to identify a set of paired interactions (“hits”) that were significantly different from WT by both ANC and CNA (Smith et al., 2016). ANC identified statistically significant differences in bead-fluorescence distributions in paired samples from WT and mutant mice by repeated two-sample Kolomogorov-Smirnov (KS) tests across experimental replications. Batch effects were removed using the ComBat function in the RStudio weighted gene correlation network analysis (v. 3.4.1; Johnson et al., 2007; Langfelder and Horvath, 2008). Significant change required differences to be present in at least 3 of 4 paired samples at *p* < 0.05 after outlier and Bonferroni-style correction. CNA identified co-varying modules of protein interaction due to *Shank3* mutation. Interactions with median fluorescence intensities (MFI) at the noise floor (MFI < 100) were not considered in CNA. The MFI of each fluorescence distribution was averaged across 4 experimental replications and input into the gene correlation network analysis. Soft-thresholding using a power adjacency function determined the power value resulting in the best approximation of scale-free topology. The minimum module size was set between 5 and 10 and typically produced 5 to 10 modules per analysis. Modules with eigenvectors significantly (*p* < 0.05) correlated with change due to *Shank3* mutation were identified and protein interactions belonging to those modules were included when the error probability of membership was below 5%. For protein pairs with significant change resulting from different epitope combinations, the measurement with the greatest change was presented. Interactome plots were made with Cytoscape (v. 3.6.1).

### Principal component analysis (PCA)

PCA was performed on ComBat, log2-transformed QMI data using all pairwise changes in protein-association as independent variables (SYSTAT, v. 13). Interactions with median fluorescence intensities (MFI) at the noise floor (MFI < 100) were not considered. The positions of individual mice in separate analyses of ShankB and ShankC mutants were plotted in Euclidian space defined by the first 2 principal components. Raw log2-fold change values were plotted as a function each protein-association’s squared component loading to indicate the weight of every association’s contribution to PCA separation. Regions of significant deviation in log2-fold change were identified by two-tailed Z-test calculated using the mean and standard deviation of all PiSCES. Associations within a region of significant Z-score change were tested for deviation from zero by unpaired t-test and those showing significant change were plotted.

### Electrophysiology

Completely-randomized and mixed-effects analysis of variance (ANOVA) were followed by Dunnett’s *post-hoc* test after a significant F statistic. Analysis of covariance (ANCOVA) examined effects of NMDAR activation on electrical coupling, adjusting for covarying strengths of baseline coupling. One and two sample t-tests were used as noted. Significance was *p* < 0.05. Data are presented as the mean ± standard error of the mean (SEM).

## SUPPLEMENTAL INFORMATION

### IP probe antibody panels used for QMI

#### Scaffold and structural proteins

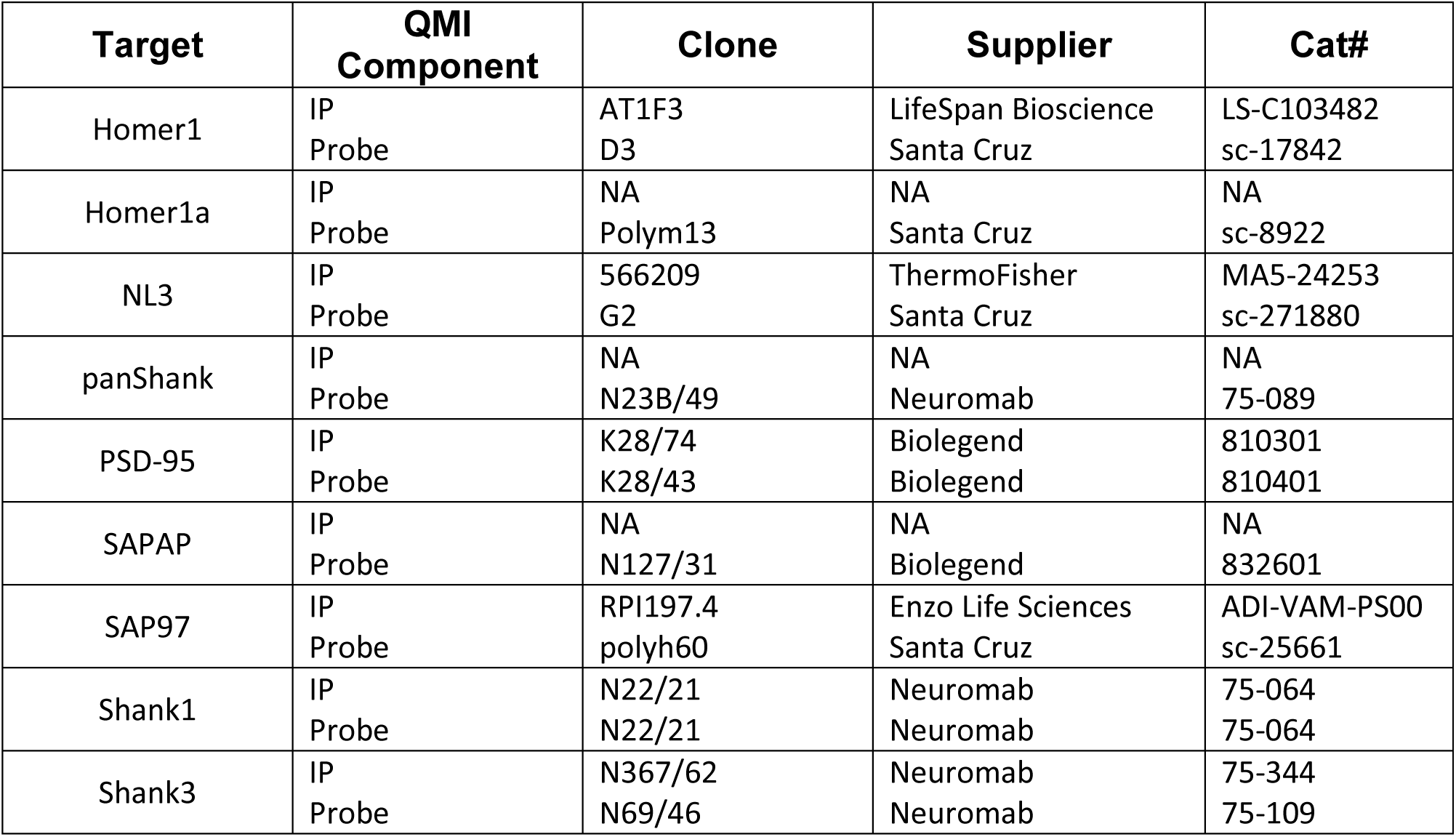

#### Receptor proteins and Cx36

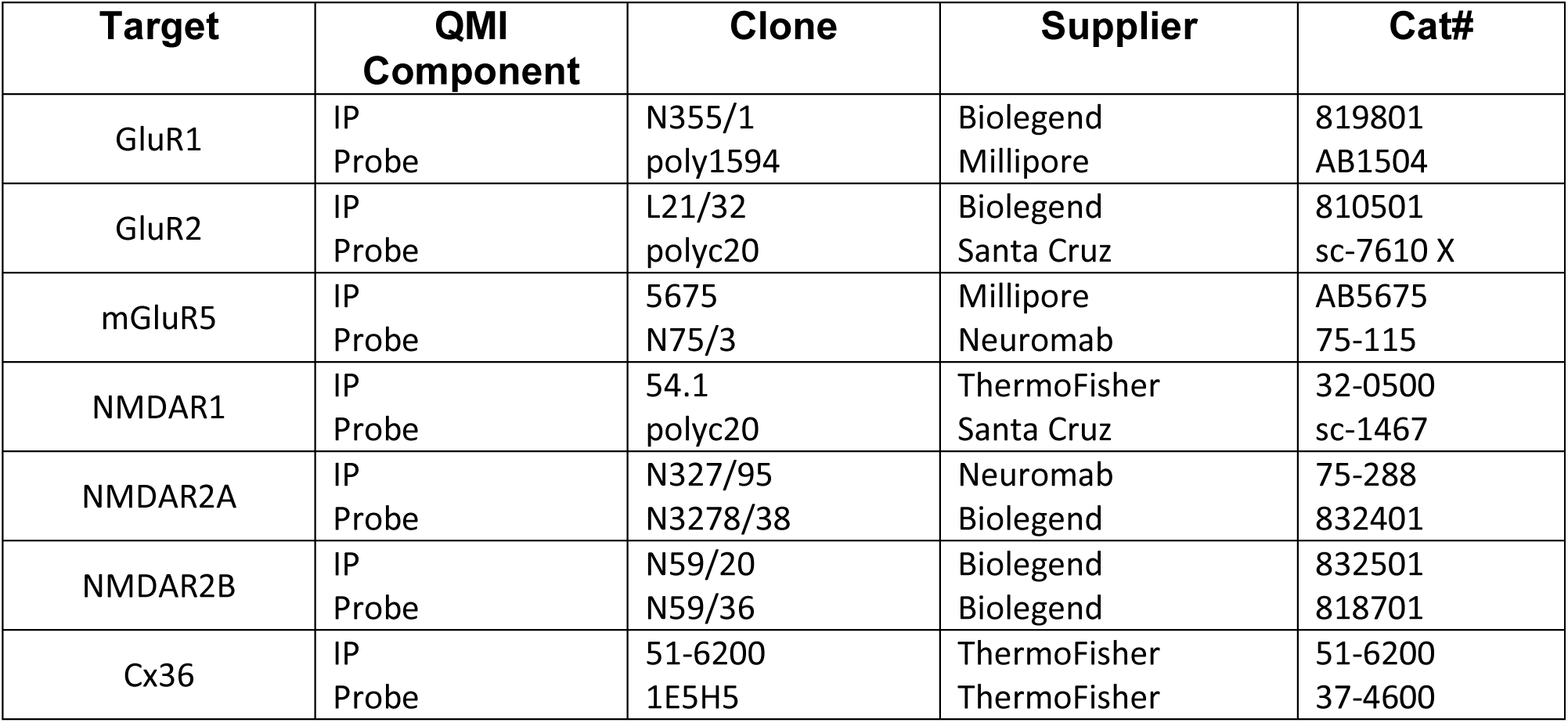

#### Signaling proteins

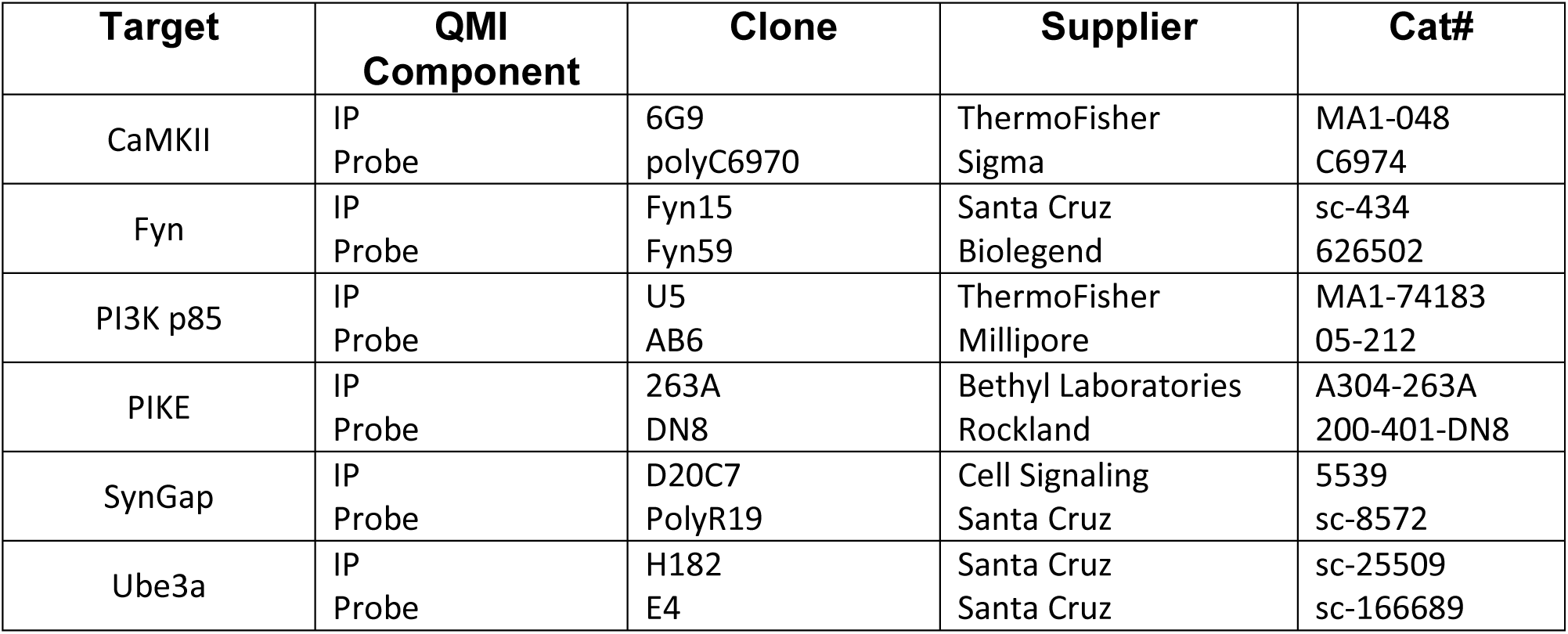

| Target | QMI Component | Clone | Supplier | Cat# |
| --- | --- | --- | --- | --- |
| CaMKII | IP<br>Probe | 6G9<br>polyC6970 | ThermoFisher<br>Sigma | MA1-048<br>C6974 |
| Fyn | IP<br>Probe | Fyn15<br>Fyn59 | Santa Cruz<br>Biolegend | sc-434<br>626502 |
| PI3K p85 | IP<br>Probe | U5<br>AB6 | ThermoFisher<br>Millipore | MA1-74183<br>05-212 |
| PIKE | IP<br>Probe | 263A<br>DN8 | Bethyl Laboratories<br>Rockland | A304-263A<br>200-401-DN8 |
| SynGap | IP<br>Probe | D20C7<br>PolyR19 | Cell Signaling<br>Santa Cruz | 5539<br>sc-8572 |
| Ube3a | IP<br>Probe | H182<br>E4 | Santa Cruz<br>Santa Cruz | sc-25509<br>sc-166689 |

Further description of the QMI method can be seen at: https://www.jove.com/t/60029/quantification-protein-interaction-network-dynamics-using-multiplexed

